# Structure of a proton-powered molecular motor that drives protein transport and gliding motility

**DOI:** 10.1101/2020.05.11.089193

**Authors:** Rory Hennell James, Justin C. Deme, Andreas Kjӕr, Felicity Alcock, Augustinas Silale, Frédéric Lauber, Ben C. Berks, Susan M. Lea

**Author notes:** These authors contributed equally to this work (Rory Hennell James, Justin C. Deme). To whom correspondence should be addressed. S.M.Lea; B.C.Berks.

## Abstract

Ion-driven motors are rare in biology. The archetypes of the three classes identified to date are ATP synthase, the bacterial flagellar motor, and a proton-driven motor that powers gliding motility and protein secretion in *Bacteroidetes* bacteria. Whilst the molecular mechanism of ATP synthase is now well understood, structural information is lacking for the other two classes of motor. Here we present the structure of the *Bacteroidetes* gliding motility motor determined by cryo-electron microscopy. The motor is an asymmetric inner membrane protein complex in which the single transmembrane helices of two periplasm-spanning GldM proteins are positioned within a ring of five GldL proteins. Combining mutagenesis and single-molecule tracking, we identify protonatable amino acid residues within the transmembrane domain of the complex that are important for motor function. Our data imply a mechanism in which proton flow leads the periplasm-spanning GldM dimer to rotate with respect to the intra-membrane GldL ring to drive processes at the bacterial outer membrane. This work provides a molecular basis for understanding how the gliding motility motor is able to transduce the energy of the inner membrane protonmotive force across the bacterial cell envelope.

## Main

The use of transmembrane ion-motive electrochemical gradients to drive biochemical processes is one of the fundamental features of cellular life^1^. Most commonly ion movement down the electrochemical gradient is coupled to transport processes or signalling. However, ion gradients can also be exploited to power the mechanical motions of membrane-associated molecular motors. Three classes of ion-driven machines are known. In ATP synthases the ion-motive gradient drives a rotary movement that results in the energy-requiring release of ATP from the catalytic sites of the enzyme^2,3^. A second class of ion-driven mechanical motor is associated with the bacterial flagellum that drives swimming motility. In this system, ion flow through stator units linked to the cell wall results in a mechanical force on the base of the flagellum that causes the flagellum to rotate^4,5^(Deme et al., companion paper). Additional members of the flagellar stator family include the periplasm-spanning Ton and Tol complexes which mechanically drive processes at the outer membrane (OM) of Gram-negative bacteria using the inner membrane (IM) protonmotive force (PMF)^6,7^(Deme et al. companion paper). The third type of ion driven motor is found in bacteria of the phylum *Bacteroidetes.* The motor in these bacteria powers rapid gliding motility across solid surfaces^8^ using the PMF across the IM as the energy source ^9–11^. The gliding motility motor generates a rotary motion at the cell surface ^11^ that results in a helical flow of surface adhesins ^9,12,13^, possibly by driving a mobile track to which the adhesins are attached ^14^.

Work with the model gliding bacterium *Flavobacterium johnsoniae* strongly suggests that the two conserved IM proteins GldL and GldM form the PMF-transducing gliding motility motor^15^(Fig. 1a). Neither of these proteins has detectable sequence similarity to the components of the other classes of ion-driven motors. GldL is composed of two predicted transmembrane helices (TMH) followed by a long cytoplasmic tail ^16^. GldM is mainly periplasmic and anchored to the IM by a N-terminal TMH ^16^. A recent structure of the periplasmic portion of GldM reveals that it contains four folded domains which assemble as a homodimeric rod that is long enough (180Å) to span the periplasm^17^. The distal end of GldM contacts a large ring structure at the OM, composed of the proteins GldK and GldN, which is thought to link the GldLM motor to the gliding apparatus ^17,18^.

**Fig. 1.**
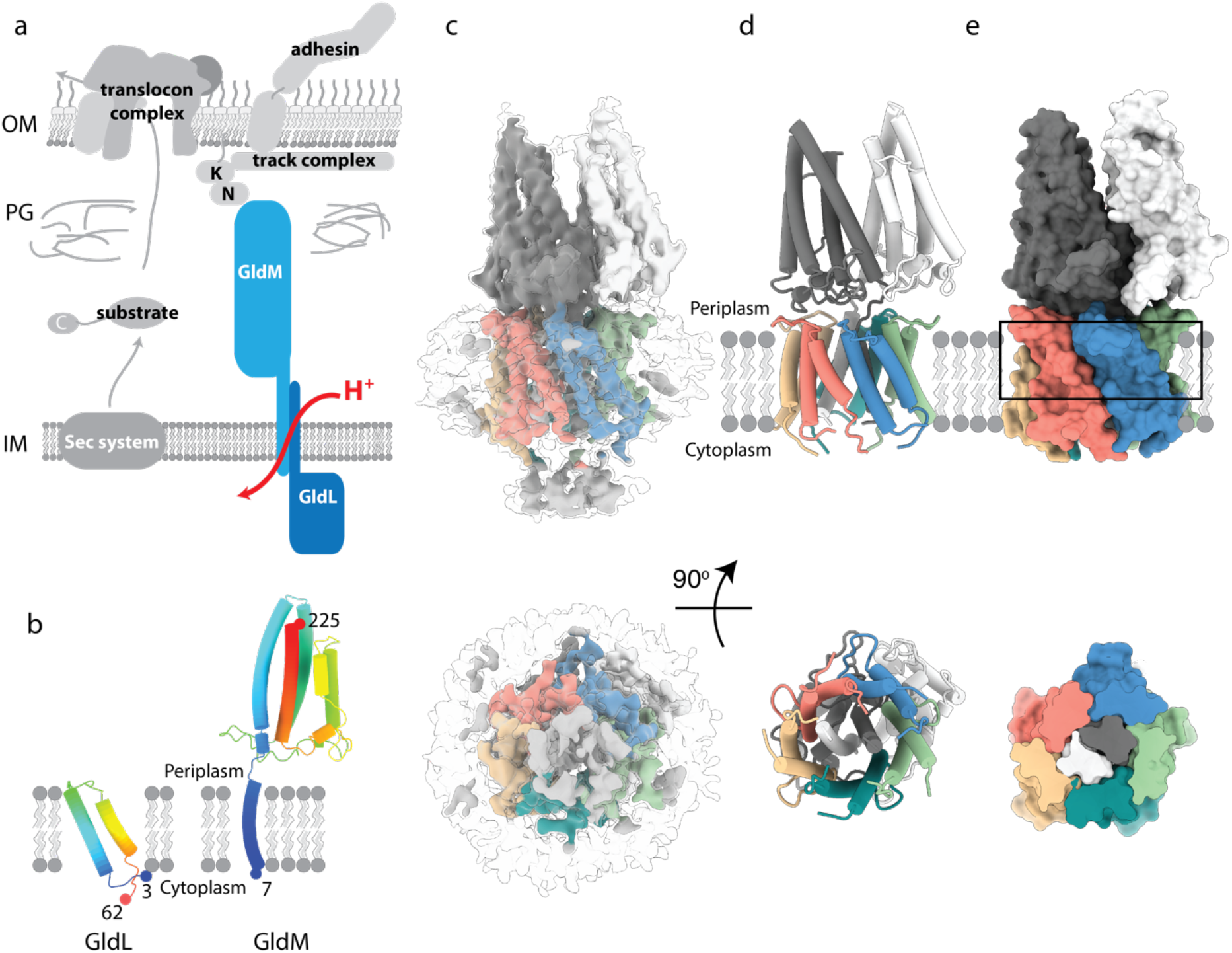
Structure of the *F. johnsoniae* GldLM’ complex. **a**, Schematic showing the relationship of the GldLM complex to the major components of the T9SS (left) and gliding motility (right) systems. OM, outer membrane; IM inner membrane; PG peptidoglycan. **b**, Structures of individual GldL and GldM subunits extracted from the complex. **c**, 3D cryo-EM reconstruction of the LMNG-solubilised GldLM’ complex at high (coloured by protein chain) and low (semi-transparent grey) contours. The detergent micelle is seen at the low contour level and is used to estimate the location of the membrane bilayer shown in **d**,**e**. **d**,**e** Cartoon and space-filling representations of the GldLM’ complex. The GldL subunits are coloured salmon, blue, green, teal, and yellow, and the GldM subunits are coloured white and dark grey. In **e** the lower panel shows a slab through the region indicated by the box in the upper panel. In **c**-**e** the lower panels show the complex viewed from the cytoplasm.

In addition to their role in gliding motility, GldL and GldM have been found to be essential for protein export across the OM by the *Bacteroidetes*-specific Type IX secretion system (T9SS) ^15,19^. Thus, the gliding motility motor is likely to also be involved in energizing protein movement through the T9SS apparatus^20–22^. Notably, GldLM homologues are still required for protein secretion in non-gliding *Bacteroidetes* species with a T9SS, such as *Porphyromonas gingivalis*, the causative agent of severe periodontal disease^19,22^. The involvement of GldLM in the T9SS leads to the prediction that Type 9 protein transport should be energized by the PMF. In agreement with this hypothesis, we find that protein export by the T9SS is PMF-dependent (Extended Data Fig. 1).

### The GldLM is an inherently asymmetric 5:2 subunit complex

The intact GldLM motor complex proved unsuitable for high resolution structure determination by cryo-electron microscopy because the anisotropic shape of the molecule led to a poor distribution of views. However, imaging a *F. johnsoniae* GldLM complex in which the GldM protein has been C-terminally truncated after the first folded periplasmic domain (hereafter referred to as GldLM’) allowed the calculation of an EM volume at 3.9 Å resolution and *de novo* building of a full atomic model (Fig. 1b-e, Extended Data Table 1, Extended Data Fig. 2-3). The transmembrane domain of the complex is fully defined in the resulting model (Fig. 1c-e). GldL is ordered between residue 3 and 62 (Fig 1b) with 153 residues of the cytoplasmic C-terminus not resolved in the current volume. The ordered portion of GldL forms a pair of TMHs arranged at ~25° to the membrane normal with a well-structured loop joining them on the periplasmic face of the membrane (Fig. 1b). Almost the entirety of the truncated GldM construct could be modelled (residue 7 to 225, Fig. 1b). The single-pass TMH is resolved and is topped by the periplasmically-located helical domain (0.7Å RMSD to the X-ray structure) (Fig. 1b). The overall complex is formed from five copies of GldL and two copies of GldM (Fig. 1d,e) with the GldL TMH pairs forming a distorted pentameric cage enclosing the two copies of the GldM TMH. Consequently, the predicted TMH of GldM is found entirely within a proteinaceous environment with no exposure to the lipid bilayer (Fig. 1e). The periplasmic domains of the GldM subunits pack against the top of the ring of GldL TMHs (Fig. 1d,e).

The overall structure of the GldLM’ complex is strikingly asymmetric both in the plane of the membrane, due to the stoichiometry mismatch of the two types of subunit within the TM helix bundle, and because the periplasmic domains of GldM adopt different tilts relative to the top of their TMHs (Fig. 1d,e). The result is that both within the membrane domain, and at the periplasmic subunit interface, each of the five copies of GldL make different contacts to GldM (Extended Data Table 2). Notably, only a single conformation of the GldLM’ complex is seen in our current data. Within the GldLM’ complex the TMHs are closely packed (Fig. 1e), implying that conformational change in the TMH of one subunit of the complex will only be possible if there are concerted conformational changes in the other subunits. As expected, the surfaces buried between the subunits of the GldLM’ complex are highly conserved and the exposed surfaces highly variable (Extended Data Fig. 4).

### Identification of functionally important residues within the transmembrane domain of the GldLM complex

The membrane domain of a proton-driven motor is anticipated to contain protonatable amino acid side chains that function to couple the transmembrane proton flow to the conformational changes required to do mechanical work. We, therefore, used site directed amino acid substitutions to test the functional importance of polar residues within the GldLM transmembrane helical bundle.

No GldM protein was detected for variants with non-conservative substitutions of GldM_R9_, probably because this residue functions as a topogenic signal for membrane protein insertion ^23^(Extended Data Fig. 5). The GldL_E49D_ variant also had reduced levels of protein expression, but this change is insufficient to account for the complete null phenotype of this variant (below).

Gliding motility was assessed by spreading behaviour on agar plates (Fig. 2a) and by microscopic examination of gliding on glass (Extended Data Table 3 and Supplementary Video 1). T9SS activity in the mutant strains was assessed through monitoring secreted chitinase activity (Fig. 2b). The effects of the substitutions on gliding were more severe than on T9SS function, suggesting that motility requires a higher level of motor function. The gliding phenotypes of the GldLM variants will reflect not only the direct effect of the substitutions on adhesin movement but also their influence on the T9SS-dependent export of the adhesins^15^. To assess the effects of the motor protein substitutions on adhesin flow independent of T9SS function, we tracked the movements of individual fluorophore-labelled adhesin molecules. In mutants that showed impaired gliding, but which retained the ability to export adhesins to the cell surface, the adhesin molecules were still observed to move in helical patterns. However, the average speed of adhesin movement was dramatically slower than in the parental strain (Fig. 2c,d and Supplementary Videos 2-7). Thus, the GldLM substitutions in these strains directly affect the mechanical force driving adhesin movement, as expected of defects in the gliding motility motor.

**Fig. 2.**
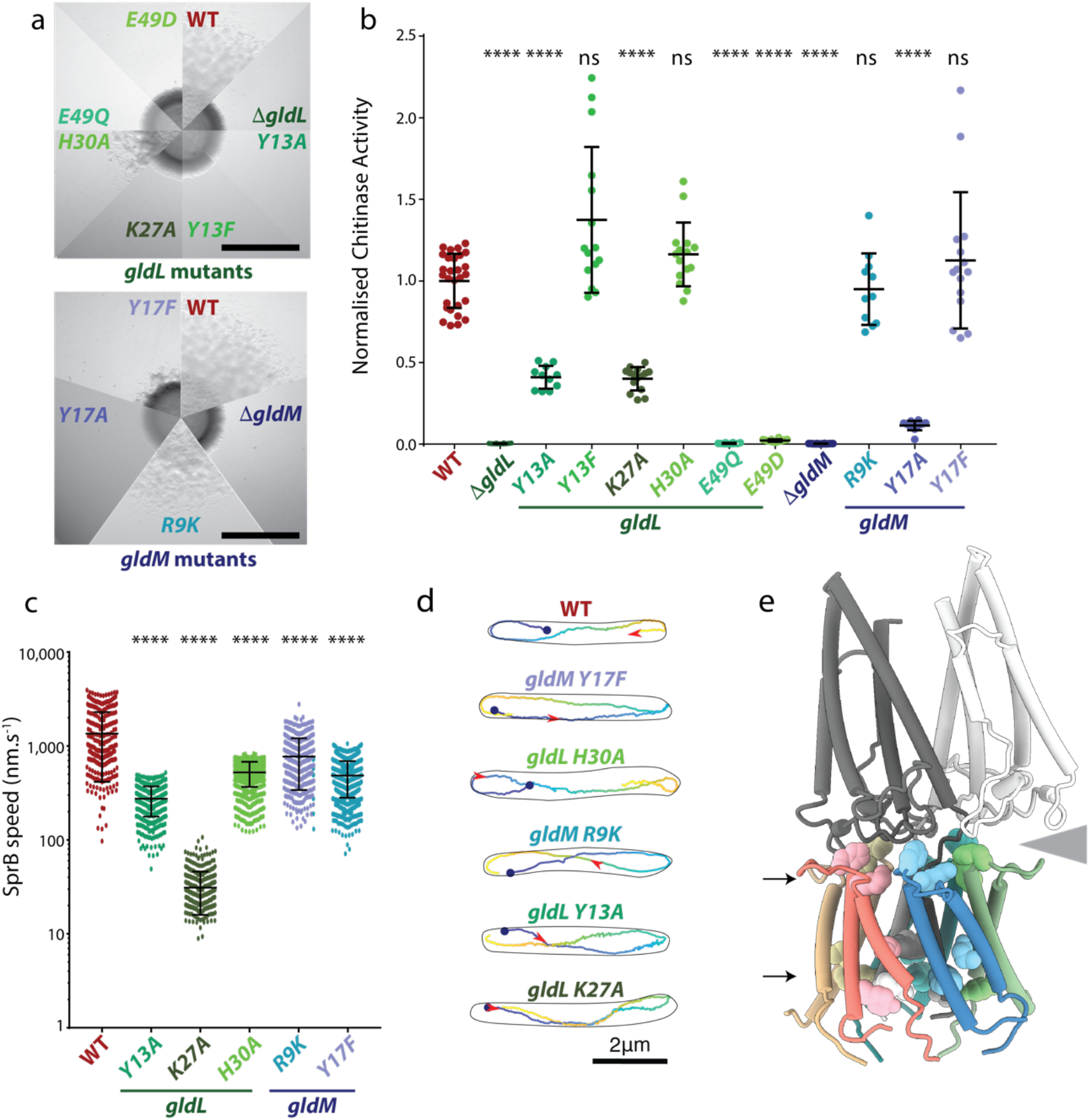
Functional analysis of protonatable residues in the transmembrane domain of GldLM. **a**-**d**, Phenotypic comparisons between wild type (WT) cells and *gldLM* single codon mutants. **a**, Gliding motility assessed by spreading on agar plates. **b**, Extracellular levels of the T9SS-secreted enzyme chitinase. Error bars show 1 standard deviation. Significance of differences relative to WT were assessed with Dunn’s multiple comparisons test (non normally distributed) or Dunnet’s multiple comparison test. **** = p < 0.0001, ns = not significant. At least 3 technical replicates of at least 3 biological repeats were performed for each strain. **c**-**d**, Cell surface movement of single fluorophore-labelled SprB adhesin molecules. **c**, Plot showing the speed of helically moving adhesins in a WT (321 tracks), *gldL*_*Y13A*_ (499 tracks), *gldL*_*K27A*_ (369 tracks), *gldL*_*H30A*_ (2,259 tracks), *gldM*_*R9K*_ (429 tracks)or *gldM*_*Y17F*_ (1519 tracks) background. For each strain the tracks are pooled data from the analysis of 3 independent cultures. Error bars show 1 standard deviation. Significance of differences relative to the WT was assessed with Dunn’s multiple comparisons test. **** = p < 0.0001. **d**, Examples of adhesin helical tracks extracted from Supplementary Videos 2-7 and coloured from blue (start) to yellow (end). The red arrowhead marks the position along the track that the adhesin has reached at 3.5s after the start point (blue dot). **e**, Protonatable residues in GldLM that exhibit reduced function when substituted. The residues are show in space-filling representation on a cartoon model of GldLM’ and fall in two bands (indicated with arrows) located at the periplasmic GldL-GldM interface and close to the cytoplasmic end of the TMH bundle. The grey wedge indicates the space between one of the GldM subunits and the top of the GldL ring.

The results of our amino acid substitution experiments (Extended Data Table 3) show that functionally important protonatable amino acids are clustered both within the membrane core of the GldLM complex and at the periplasmic GldL/GldM interface (Fig. 2e). The tight packing of the transmembrane helix bundle (Fig. 1e) seen in the structure provides no route for proton movement between these two layers of residues. This implies that the motor cycle must include a conformational change to open an aqueous channel between these regions at one point in the ring.

Within the membrane core the invariant residues GldL_E49_ and GldM_Y17_ and the highly conserved residues GldL_Y13_ and GldM_R9_ are important for motor function. The symmetry mis-match in the GldLM’ complex means that within each chain these residues are in a different environment. Thus, in one GldM copy GldM_R9_ (chain B; coloured grey in the figures) is positioned to form a salt bridge with GldL_E49_ (chain E, coloured salmon in the figures), whilst GldM_Y17_ (chain A, coloured white in the figures) and GldL_Y13_ (chain E) bracket this pair of residues (Fig 3a). By contrast, GldM_R9_ in the other copy of GldM (chain A) is rotated away from GldL_E49_ on the nearest GldL subunit and so these two residues are no longer positioned to interact. The substitution data show that protonation of the GldL_Y13_ and GldM_Y17_ side chains is important for their function, as expected if these residues are involved in transducing transmembrane proton movements to mechanical work. However, the substitution data also show the importance of the aromatic nature of GldL_Y13_ and GldM_Y17_, suggesting that these residues are additionally involved in mechanistically important packing interactions.

**Fig. 3.**
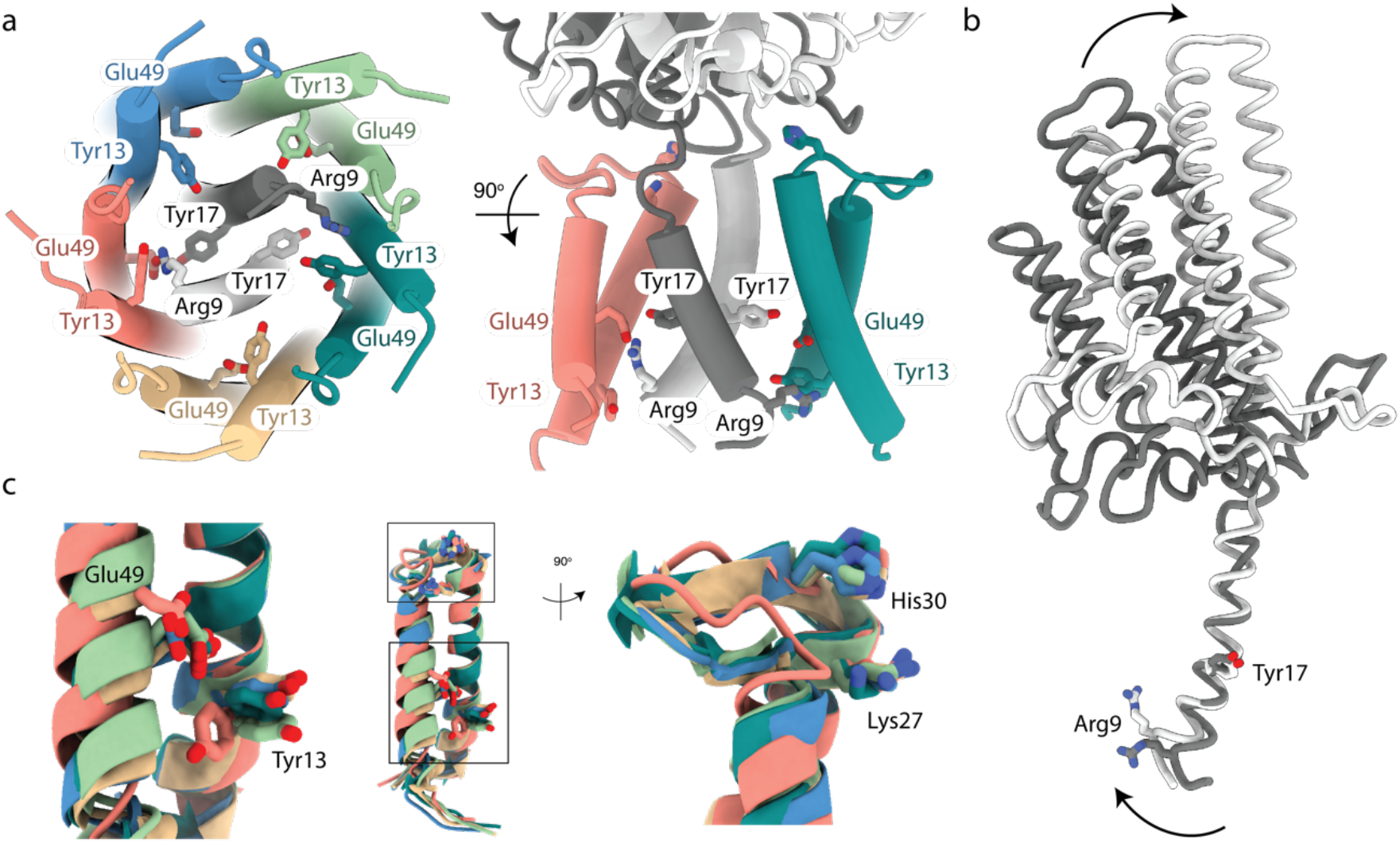
The structural asymmetry of the GldLM’ complex leads to important differences in residue environments between chains. The side chains of functionally important resides are shown in stick representation with oxygen atoms red and nitrogen atoms blue. Proteins chains are coloured as in Fig. 1 c-e. **a**, The side chains of functionally important resides near the cytoplasmic end of the TMH bundle. For clarity, only two GldL subunits are shown in the right hand view. **b**, Overlay of the two copies of GldM by superposition of the TMH. **c**, Superposition of the five copies of GldL by overlaying TMH2. The boxes in the central panel indicating the positions of the two regions magnified on either side.

### Structural asymmetry in the GldLM complex

An overlay of the two GldM’ subunits shows that the cytoplasmic end of the TMH must be bent in order for GldM_R9_ to ion pair with GldL_E49_ (Fig. 3b). In this ion-paired copy of GldM the angle between the TMH and the periplasmic domain is less acute than in the other GldM chain, and so the periplasmic domain is less tilted relative to the GldL ring (Figs. 2e and 3c). Indeed, it is the periplasmic domain of the GldM subunit that is not participating in the ion pair that tilts sufficiently to make extensive interactions with the top of the ion-paired GldL subunit (Fig. 2e and 3b,c).

An overlay of the five copies of the GldL protein found in the GldLM’ complex reveals that the helix in which GldL_E49_ is positioned to ion pair with GldM_R9_ is shifted compared to the equivalent helix in the other copies of GldL (Fig. 3c). This helix movement is accompanied by a reorientation of GldL_Y13_ located on the second TMH of the same chain and a remodelling of the periplasmic loop that is packed against the GldM periplasmic domain (Fig. 3c). These changes in GldL conformation are likely linked to conformational differences between the two copies of GldM and imply that mechanical forces exerted on the GldM subunits within the membrane are reinforced by interactions between the GldM periplasmic domains and the top of the GldL ring. Notably, substitution of two invariant residues (GldL_K17_ and GldL_H20_) within the GldL periplasmic loop that appear to mediate interactions with the GldM periplasmic domain significantly impair gliding motility (Figs. 2a,c,d and 3c).

Taken together these observations show that the presence or absence of the GldM_R9_ – GldL_E49_ ion pair in the two copies of the GldM protein is linked to conformational differences between the proteins. This suggests a simple model of reciprocal conformational change between the two GldM copies to produce mechanical force in the periplasm. In this model the GldM_R9_ – GldL_E49_ ion pair involving one GldM subunit is broken by protonation of GldL_E49_ by an incoming proton from the periplasmic side of the membrane, subsequent to which GldM_R9_ on the other GldM subunit forms an equivalent ion pair through ejection of a proton from GldL_E49_ on the nearest GldL molecule. GldM_Y17_ and GldL_Y13_ are well-positioned to act as the proton donor and acceptor, respectively, to the GldM_R9_ – GldL_E49_ ion pair. In this context, the movement of the GldL_Y13_ side chain away from the ion pair in the salt-bridged copy of GldL might be a mechanism to prevent the immediate short-circuiting transfer of the proton to the cytoplasm following protonation of the ion pair. Co-operative movements of the two GldM proteins will be reinforced by the fact that it is the GldM subunit that is not involved in the ion pair that both provides the apparent proton donor to the ion pair (through GldM_Y17_) and that possesses the periplasmic domain that forms intimate contacts with the periplasmic loops of the GldL subunit involved in the ion pair.

### Orientation of the GldM periplasmic domains

An overlay of the first periplasmic domains of GldM in our cryoEM structure with the equivalent domains in the previously reported crystal structures of the dimeric, isolated, GldM periplasmic region^17^ demonstrates that the two copies of the domain are splayed apart in the cryoEM structure compared to the earlier crystal structure (Fig. 4a). We were concerned that this difference between the two structures might have arisen due to our truncation of the periplasmic region of GldM, even though neither co-variance nor conservation suggest a strong dimerization interface between these first periplasmic domains (Extended Data Fig. 4) and the isolated domain from the *P. gingivalis* homologue crystallises as a monomer^17^. To resolve this issue we turned to lower resolution cryoEM volumes derived from images of full length GldLM complexes. Data from the *P. gingivalis* GldLM homologue PorLM were of sufficient quality to allow location of the two TMH helices of PorM enabling us to position the atomic model for *F. johnsoniae* GldLM’ within the lower resolution *P. gingivalis* PorLM volume (Fig 4b,c). Interpretation of the lower resolution volume in this way demonstrates that even in the context of the full length GldM homologue the GldM first periplasmic domains are splayed with an acute (and different between the two copies) angle between the first and second domains (Fig 4c). Density for the second periplasmic domain is also visible in the volume and is compatible with the strand-swapped dimeric arrangement of this domain seen in the crystal lattice despite the separation of the first domain. The splaying of GldM is likely driven by packing of the first periplasmic domains onto the top of the GldL transmembrane domains and packing of the N-terminal helices within the GldL cage. The need to anneal the two GldM chains above this domain may explain the unusual strand-swapped dimer that forms the next (second) periplasmic domain. More distal domains were only present in the volume at very low contour levels presumably due both to their distance from the centre of the alignment and to some conformational variability of the entire periplasmic portion with respect to the transmembrane region. Nevertheless, no further large deviations from a linear conformation are seen for the distal domains. Combining our structural information with the earlier crystallographic data generates a composite model for the GldLM complex in which the rod-like periplasmic region of GldM projects into the periplasm at an appreciable angle away from the membrane normal (Fig. 4d).

**Fig. 4.**
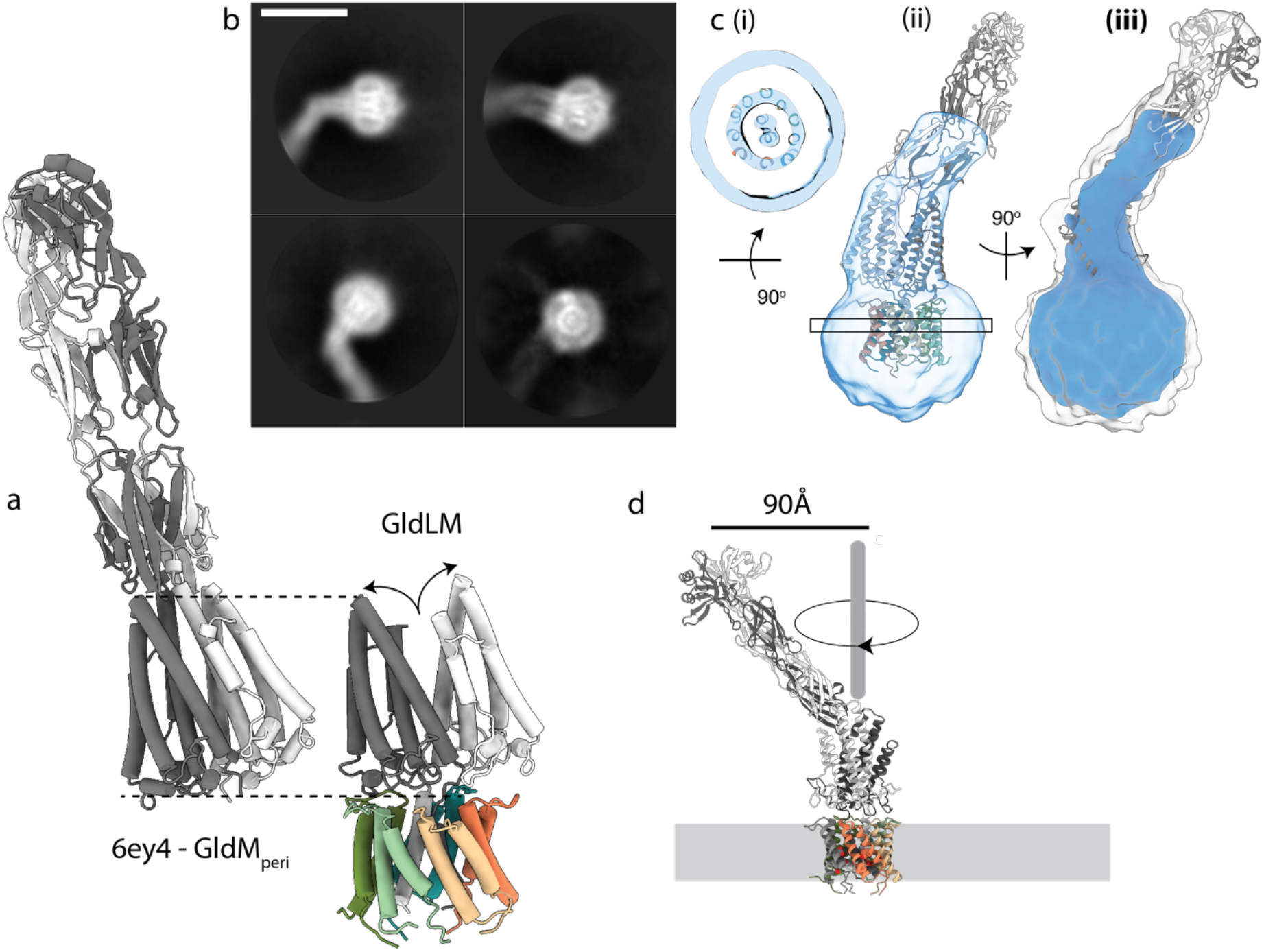
Structure of the full length motor complex. **a**, Comparison of the crystal structure of the periplasmic domains of GldM (PDB 6ey4, GldM_peri_) with the cryoEM structure of the periplasmically truncated GldLM’ complex. The structures are displayed with the first periplasmic domain of the dark grey GldM chain aligned (dotted lines) and demonstrate the splaying (curved arrows) of the GldM chains in the context of the truncated complex. **b**, Representative 2D classes from micrographs of the full-length *P. gingivalis* motor (PorLM) showing detail in the transmembrane region (all 4 panels), a large bend between the first two periplasmic domains of PorM (left panels), and splaying of the first periplasmic domains of the two copies of PorM (upper right panel). Scale bar 100 Å. **c**, Different views of the *P. gingivalis* motor 3D cryo-EM volume at a high (blue) and low (white) contour level. (**i**) The PorM TMHs can be resolved allowing confident placement of the *F. johnsoniae* GldLM’ model in the volume. (**ii**) Splaying of the PorM first periplasmic domain pair is seen as in the *F. johnsoniae* GldLM’ complex. (**jjj**) The extension to the volume seen at a low contour level matches well to the length and shape of the dimeric GldM periplasmic domains 2-4 seen in the crystal structure. **d**, Cartoon to demonstrate how the bend between periplasmic domains 1 and 2 would allow the tip of a rotating GldM/PorM to define a large track at the outer-membrane.

## Discussion

The GldLM’ structure reveals an asymmetric complex bedded in the inner membrane. The lack of shared symmetry between the components of the GldLM’ complex, together with the caging of GldM within GldL, suggests that rotation of GldM within the GldL ring is the most likely mechanical movement produced by the motor. This type of motion would be consistent with the observation that the gliding motility system is able to generate rotary motion at the cell surface^11^. In order for the GldLM motor to drive processes at the OM through rotation of the distal end of GldM, GldL must remain stationary in the IM to act as a stator to the rotating GldM protein. GldL cannot be immobilized by binding to the cell wall since GldM covers the parts of GldL that are exposed at the periplasmic side of the membrane. Instead, it is likely that GldL is held in position through interactions between the currently unvisualized cytoplasmic tail and a static cytoplasmic structure. Sequential morphing of the structure between the GldL conformational states was used to produce an animation that indicates the approximate molecular changes that would occur in the GldLM complex on rotation of the GldM periplasmic domain (Supplementary Video 8).

The arrangement of the ten GldL TMHs around two central GldM TMHs is reminiscent of the structural organization of the transmembrane core of the flagellar MotAB stator complex (reported in our companion paper (Deme et al. companion paper)) and the related ExbBD complex of the Ton system ^6^(Deme et al. companion paper) even though the mobile and stationary components of the GldLM motor are reversed relative to MotAB. This organizational similarity is particularly remarkable given the lack of detectable sequence similarity between GldLM and the subunits of the other motor complexes. The implications of this protein architecture being shared between bacterial ion-driven motors are discussed in the companion paper (Deme et al., companion paper).

## Acknowledgements

We thank Mark McBride and Yongtao Zhu for providing reagents for the genetic manipulation of *F. johnsoniae*. We thank Luke Lavis for supplying the Janelia Fluor ligands, Sam Hickman for advice on fluorescence imaging, and Alan Wainman for assistance with phase contrast imaging. We thank Paul Guy for his involvement in producing the pulse-chase mCherry-CTD fusion construct and Richard Berry for discussions and comments on the manuscript. We acknowledge the use of the Central Oxford Structural Microscopy and Imaging Centre (COSMIC), the Oxford Micron Advanced Imaging Facility, and the Oxford Advanced Proteomics Facility. We thank Kevin Foster for providing additional imaging facilities. This work was supported by Wellcome Trust studentships to R.H.J. and A.S., a Biotechnology and Biological Sciences Research Council studentship to A.K., and Wellcome Trust Investigator Awards 107929/Z/15/Z and 100298/Z/12/Z. COSMIC was supported by a Wellcome Trust Collaborative Award 201536/Z/16/Z, the Wolfson Foundation, a Royal Society Wolfson Refurbishment Grant, the John Fell Fund, and the EPA and Cephalosporin Trusts.

## Author contributions

R.H.J carried out all genetic and biochemical work except as credited otherwise. J.C.D. and S.M.L. collected EM data and determined the structure. A.K. developed and carried out the SprB tracking experiments including strain construction. F.A. carried out the pulse chase analysis of protein export, A.S. assayed cellular ATP levels, F.L. constructed the Δ*gldL* strain and produced the GldL and GldM antibodies. B.C.B. and S.M.L. conceived the project. All authors interpreted data and wrote the manuscript.

## Methods

No statistical methods were used to predetermine sample size. The experiments were not randomized and the investigators were not blinded to allocation during experiments and outcome assessment.

### Bacterial strains and growth conditions

All strains and plasmids used in this work are listed in Extended Data Tables 4 and 5. *F. johnsoniae* was routinely grown in Casitone Yeast Extract (CYE) medium^24^ at 30 °C with shaking. To assess motility on glass and for chitinase secretion measurements cells were grown in Motility Medium^25^ (MM). PY2 medium^26^ was used to assess motility on agar plates. *E. coli* cells were routinely grown in Luria Bertani (LB) medium at 37 °C with shaking. When required, antibiotics were added at the following concentrations: erythromycin, 100 μg ml^−1^; spectinomycin, 100 μg ml^−1^; ampicillin, 100 μg ml^−1^; kanamycin, 30 μg ml^−1^.

### Genetic constructs

All primers used in this work are described in Supplementary Data Table 1. All plasmid constructs were verified by sequencing.

A vector directing the co-expression of *F. johnsoniae* GldL and C-terminally truncated GldM was produced as follows. The chromosomal region encoding GldL and the first 225 amino acids of GldM was amplified using primers RHJ164 and RHJ165. The plasmid pT12^27^ was linearized by amplification with primers RHJ162 and RHJ163. The two fragments were assembled by Gibson cloning to yield plasmid pRHJ007. A vector for expression of *F. johnsoniae* GldL and the first 232 amino acids of GldM was produced similarly, using primer pair RHJ166 & RHJ167 to create pRHJ008.

A vector directing the co-expression of *P. gingivalis* ATCC 33277 PorL and PorM in *E. coli* was produced as follows. pWALDO-sfGFP^28^ was digested with BamHI and HindIII and the resulting fragment coding for the TEV cleavage site, superfolder GFP, and a His_8_ tag was ligated into the corresponding sites in the first multiple cloning site of pCDFDuet-1 (Novagen), yielding plasmid pRHJ001. A C-terminal Twin-Strep tag-coding sequence was added to the second multi-cloning site of pRHJ001 by Q5 site-directed mutagenesis (New England Biolabs) using primers RHJ025 and RHJ026, giving plasmid pRHJ002. The NcoI site in the superfolder GFP-coding sequence was removed by QuikChange mutagenesis (Agilent) using primers RHJ046 and RHJ047, yielding plasmid pRHJ003. pRHJ003 was digested with NcoI and XhoI and the resulting fragment ligated into the corresponding sites of pETDuet-1 to give pRHJ004. *P. gingivalis porL* was amplified from genomic DNA with primers RHJ051 and RHJ052. The resulting 0.9 kb fragment was inserted between the NcoI and BamHI sites of pRHJ004 to give pRHJ005. *P. gingivalis porM* was amplified with primers RHJ057 and RHJ058. The resulting 1.6 kb fragment was inserted between the NdeI and KpnI sites of pRHJ005 to give pRHJ006.

A suicide vector to produce an in-frame unmarked deletion of *F. johnsoniae gldL* was produced as follows. A 2.6 kbp region corresponding to the first 36 bp of *gldL* together with the directly upstream region was amplified with primers FL309 and FL310. A 2.5 kbp region corresponding to the final 36 bp of *gldL* together with the directly downstream region was amplified with primers FL311 and FL312. The vector pYT313^29^ was linearized by amplification with primers FL313 and FL314. These three fragments were then assembled by Gibson cloning to give plasmid pFL89.

A suicide vector to produce an in-frame unmarked deletion of *F. johnsoniae gldM* was produced as follows. A 2.7 kb region corresponding to the first 31 bp of *gldM* together with the directly upstream region was amplified with primers RHJ148 and RHJ160. A 2.5 kb region corresponding to the final 38 bp of *gldM* together with the directly downstream region was amplified using primers RHJ161 and RHJ147. The vector pGEM-T was linearized by amplification using primer RHJ144 and RHJ145. These three fragments were assembled by Gibson cloning to yield pRHJ010. The fragment containing the two *F. johnsoniae* chromosomal regions was then amplified from pRHJ010 with primers RHJ110 and RHJ113. The resulting 5.2 kb fragment was inserted between the BamHI and KpnI sites of pYT354^29^ to give plasmid pRHJ011.

The strategy to generate point mutations in *gldL* involved the use of an intermediate cloning vector that was produced as follows. A 5.3 kb region including the *gldL* gene sequence and surrounding chromosomal regions was amplified using primers RHJ146 and RHJ149. pGEM-T was linearized by amplification with primers RHJ144 and RHJ145. The two fragments were assembled by Gibson cloning to give pRHJ012. An intermediate cloning vector for generating mutations in *gldM* was produced in an analogous manner using primers RHJ147 and RHJ148 and yielding plasmid pRHJ013.

A suicide vector for the introduction of the *gldL(N10A)* point mutation was produced as follows. pRHJ012 was linearized by amplification with primers RHJ310 and RHJ311, which introduce the desired 2 bp point mutation in codon N10. The resulting fragment was re-circularised by *in vivo* assembly ^30^, yielding plasmid pRHJ014. The fragment containing the mutated *gldL* sequence and adjacent regions was then amplified from pRHJ014 with primers RHJ341 and RHJ342. The vector pYT354 was linearized by digestion with BamHI and SalI. The two fragments were then assembled by *in vivo* Gibson cloning to give plasmid pRHJ036. Other point mutations were generated similarly, using pRHJ012 as a template for mutations in *gldL* and pRHJ013 as a template for mutations in *gldM*, using the primers described in Supplementary Table 1.

A suicide vector to introduce a Twin-Strep tag-coding sequence to the 3’ end of *gldL* was produced as follows. A 2.5-kbp fragment comprising *gldL* together with the directly upstream region was amplified using primers FL265 and FL266. This fragment was inserted into the SphI and NcoI sites of pGEM-T easy to generate pFL76. A 2.6-kbp fragment downstream of *gldL* was amplified using primers FL267 and FL268. This fragment was inserted into the NcoI and SalI sites of pFL76 to generate pFL77. A fragment encoding a TEV cleavage site followed by a Twin-Strep tag was amplified from pRHJ007 using primers RHJ172 and RHJ173. This fragment was inserted between the BamHI and NcoI sites of pFL77 to give plasmid pRHJ058.

A suicide vector for the construction of a *gldL(N10A)-twinstrep* strain was produced as follows. A fragment including the pGEM-T backbone, the mutation site, and a 2.3 kb region upstream of the mutation site was amplified from pRHJ012 using primers RHJ311 and RHJ144. A fragment containing the mutation site together with a 3.2 kb region downstream of the mutation site including the Twin-Strep tag-encoding sequence was amplified from pRHJ058. These fragments were assembled by Gibson cloning to give pRHJ059. The fragment containing the mutated *gldL* sequence and adjacent regions was amplified using primers RHJ341 and RHJ342. pYT354 was linearised by digestion with BamHI and SalI. These two fragments were assembled by Gibson cloning to give pRHJ081. Suicide vectors for the construction of other *gldL* point mutations in a *gldL-twinstrep* background were produced in an analogous manner using the primers described in Supplementary Table 1.

To introduce point mutations in *gldM* into a *gldL-twinstrep* background, the mutation site and the region upstream of the *gldM* point mutation, including the Twin-Strep tag-encoding sequence of the *gldL* gene, was amplified from pRHJ058. The mutation site and the region downstream of the point mutation together with the pGEM-T backbone, was amplified from pRHJ013. These fragments were assembled by Gibson cloning. Primers RHJ342 and RHJ343 were then used to transfer the mutated *gldM* sequences into pYT354.

A suicide vector for the construction of *halotag-sprB* strains was produced by Gibson assembly of the following four fragments yielding plasmid pAK021: pYT313 linearized with primers AK41 and AK62; codons 86 to 2647 of *sprB* amplified from the *F. johnsoniae* chromosome with primers AK59 and AK60; *twinstrep-halotag* amplified from plasmid pET21-ts-halo-RemA97CTD with primers AK37 and AK61; a genomic fragment extending from the start of *sprD* through to codon 87 of *sprB* was amplified from the *F. johnsoniae* chromosome with primers AK36 and AK40.

For pulse-chase experiments, a plasmid directing production of a tripartite fusion protein consisting of the *F. johnsoniae* RemA signal sequence, mCherry, and the RemA C-terminal domain under the control of the *remA* promoter was constructed as follows. A region encompassing 361 nucleotides immediately upstream of *remA* together with the first 150 nucleotides of *remA* was amplified from genomic DNA using primers PG001 and PG002, then cloned into the XbaI-SpeI restriction sites of pCP11^24^. Between the SpeI and SacI sites of the resulting plasmid was inserted the coding sequence for mCherry with no stop codon amplified from plasmid pRVCHYC-5^31^ using primers PG003 and PG004. Finally a region coding for the 97 C-terminal residues of RemA was amplified with primers PG005 and PG006 and inserted between the SacI and SalI sites to produce plasmid pCP-remA_us_-mch-CTD97_remA_. A plasmid coding for the fusion protein with a K1432A substitution in the RemA CTD was made by the QuikChange mutagenesis (Agilent) using primers PG007 and PG008.

Suicide and expression plasmids were introduced into *F. johnsoniae* strains by biparental mating using *E. coli* S17-1^32^ as the donor strain, as previously described^24^. Point mutations in *gldL* were introduced into the chromosome by using the Δ*gldL* strains Fl_082 or Ak_205 as the recipient. Point mutations in *gldM* were introduced into the chromosome by using the Δ*gldM* strains Rhj_006 or Ak_203 as the recipient. Erythromycin resistance was used to select cells with a chromosomally-integrated suicide plasmid. One of the resulting clones was grown overnight in CYE without antibiotic to allow for loss of the plasmid backbone, and then plated onto CYE agar containing 5 % sucrose. Sucrose-resistant colonies were screened by PCR for the presence of the desired chromosomal modification and then verified by sequencing.

### Purification of PorLM and GldLM complexes

GldLM’ proteins were overproduced from plasmids pRHJ007 and pRHJ008 as follows. Colonies of BL21(DE3) cells carrying the appropriate plasmid were inoculated into 50 ml 2xYT medium and cultured at 37°C with shaking for 6-8 h. The cells were diluted to OD_600_ = 0.02 in TB supplemented with 0.2 % L-rhamnose and then grown at 37°C with shaking for 14 h. Cells were harvested by centrifugation at 5,000*g* for 15 min at 4 °C. Cells were washed once in Dulbecco A phosphate buffered saline (PBS) and stored at −20°C until further use.

The frozen cell pellet was resuspended in 3.3 ml per g of cells of Lysis Buffer (PBS supplemented with 1 mM EDTA, 30 μg ml^−1^ DNase I, 400 μg ml^−1^ lysozyme, and 1 tablet per 100 ml SIGMAFAST™ protease inhibitor cocktail). The cells were then disrupted using an Emulsiflex homogeniser operated at 15,000 PSI. The resulting lysate was centrifuged at 27,000*g* for 30 min at 4 °C to remove cellular debris before the membrane fraction was recovered by centrifugation at 210,000*g* for 1 h at 4 °C. The membrane pellets were stored overnight at 4 °C before being resuspended in 8 ml Resuspension Buffer (PBS, 1 mM EDTA) per g membrane. 1 ml 10 % (w/v) lauryl maltose neopentyl glycol (LMNG; Anatrace) solution was added per g of membranes and the suspension was gently stirred at 4 °C for 2 h. Non-solubilised material was removed by centrifugation at 75,000*g* at 10 °C for 30 min. The resulting supernatant solution was passed through a 5 ml StrepTactin XT cartridge (IBA). The column was washed with 10 column volumes (CV) StrepTactin Wash Buffer (PBS, 1 mM EDTA, 0.02 % LMNG). Protein was eluted in 2 CV StrepTactin Elution Buffer (PBS, 0.02 % LMNG, 1 mM EDTA, 50 mM D-biotin). The elution fractions were concentrated to 500 μl using a 100 kDa MWCO Amicon Ultra–15 centrifugal filter, and injected on to a Superose 6 10/300 Increase GL size-exclusion chromatography column (GE Healthcare) equilibrated in 20 mM HEPES pH 7.5, 150 mM NaCl, 0.02 % LMNG. Protein purity was assessed by SDS-PAGE (Extended Data Figure 2a-b) and GldLM’-containing fractions were pooled, concentrated using a GE Healthcare 100 kDa MWCO Vivaspin 500 concentrator, and stored at 4°C until use. Protein concentrations were determined spectrophotometrically using A_280nm_ 1 = 1 mg ml^−1^.

Production of recombinant PorLM complexes was carried out as follows. An overnight culture of *E. coli* Mt56(DE3)^33^ carrying pRHJ006 in 2xYT medium^34^ was diluted 100-fold into fresh Terrific Broth medium^35^ containing 100 μg ml^−1^ ampicillin, and grown at 37°C with shaking to OD_600_ = 4.0. Protein production was then induced by addition of 0.1 mM IPTG and the cells cultured for 15 h at 24°C with shaking. Cells were harvested and proteins purified as described above for the GldLM’ complex, with the following differences. 1 mM EDTA was omitted from the StrepTactin Elution Buffer. Following elution from the StrepTactin XT cartridge, the eluted fraction was subsequently passed through a 5 ml Ni-NTA Superflow cartridge (Qiagen). The column was washed with 10 CV Ni-NTA Wash Buffer 1 (PBS, 0.02 % LMNG, 20 mM imidazole), followed by 10 CV Ni-NTA Wash Buffer 2 (PBS, 0.02 % LMNG, 40 mM imidazole). Protein was eluted in 1.2 CV Ni-NTA Elution Buffer (PBS, 0.02 % LMNG, 250 mM imidazole). 1mg of TEV-His_6_ protease was added per 10 mg eluted protein and the sample dialysed overnight against Dialysis Buffer (PBS, 0.02 % LMNG) in 3.5 kDa molecular weight cutoff (MWCO) SnakeSkin^®^ dialysis tubing (Thermo Scientific) at 4°C. The dialysate was centrifuged at 3,220*g* for 15 min at 4°C. Imidazole from a 2M stock was added to the supernatant to achieve a final concentration of 20 mM. The supernatant was passed over a Ni-NTA column. The flowthrough containing PorLM was collected and then further processed by SEC chromatography as described for the GldLM’ purification above (see Extended Data Figure 1c).

### Cryo-EM sample preparation and imaging

Four microlitre aliquots of purified GldLM’ (A_280nm_ = 0.4-0.5) or PorLM (A_280nm_ = 0.2- 0.5) complexes were applied onto glow-discharged holey carbon coated grids (Quantifoil 300 mesh, Au R1.2/1.3), adsorbed for 10 s, blotted for 2 s at 100% humidity at 4 °C and plunge frozen in liquid ethane using a Vitrobot Mark IV (FEI).

Data were collected in counting mode on a Titan Krios G3 (FEI) operating at 300 kV with a GIF energy filter (Gatan) and K2 Summit detector (Gatan) using a pixel size of 0.822 Å and a total dose of 48 e^−^/Å^2^ spread over 20 fractions.

### Cryo-EM data processing

Motion correction and dose weighting were performed using MotionCor implemented in Relion 3.0^36^. Contrast transfer functions were calculated using CTFFIND4^37^. Particles were picked in Simple^38^ and processed in Relion 3.0^36^. Gold standard Fourier shell correlations using the 0.143 criterion and local resolution estimations were calculated within Relion^36^ (Extended Data Fig. 3).

3,284,887 GldLM’ particles were extracted from 9,858 movies in 256 × 256 pixel boxes and subjected to a round of reference-free 2D classification, from which 531,515 particles were recovered. An *ab initio* initial model was generated from a subset of 500 particles using SIMPLE^38^. This model was low-pass filtered to 60 Å and used as reference for 3D classification (4 classes, 7.5° sampling) against a 335,887 particle subset followed by refinement which yielded a 6.8 Å map. This map was used as initial reference (40 Å low-pass filtered) and mask for 3D classification (3 classes, 15 iterations at 7.5° sampling then 10 iterations at 3.75° sampling) against the entire dataset and further refined to 4.0 Å. Bayesian particle polishing followed by another round of 3D classification with local angular searches yielded a 3.9 Å map from 119,230 particles after refinement.

PorLM particles (1,133,336) were extracted from 14,135 movies in 324 × 324 boxes and subjected to two rounds of reference-free 2D classification, from which 495,572 particles were recovered. After recentering and reextraction in a smaller box (256 × 256), particles were subjected to 3D classification (4 classes, 15 iterations at 7.5° sampling then 10 iterations at 3.75° sampling) against a 40 Å low-pass filtered GldLM map. Selected particles (199,929) were refined against the corresponding map (low-pass filtered to 40 Å) first using a soft spherical mask of 140 Å and then with a 180 Å mask. Recentered particles, now in a 324 × 324 pixel box, were refined using a spherical mask of 400 Å. Particle re-centering and re-extraction in 480 × 480 pixel boxes followed by refinement using local searches with a mask covering the stalk and base of PorLM yielded an 8.6 Å map.

### GldLM model building and refinement

The first periplasmic domains (residues 36-224) of the *F. johnsoniae* GldM crystal structure ^17^ (PDB 6ey4) were rigid body fitted into the GldLM’ cryo-EM density map using Coot ^39^. All other residues were built *de novo* using Coot^39^ guided by TMH predictions from TMHMM^40^. Multiple rounds of rebuilding in both the globally sharpened and local-resolution filtered maps and real-space refinement in Phenix^41^ using secondary structure, rotamer, and Ramachandran restraints yielded the final model described in Table 1. Validation was performed using Molprobity^42^. Conservation analysis was carried out using the Consurf server^43^.

The GldLM’ model and the C-terminal periplasmic domains (residues 225-515) of PorM (PDB 6ey5) were docked into the PorLM map using Chimera ^44^. Figures were prepared using UCSF ChimeraX^44^ and Pymol (The PyMOL Molecular Graphics System, Version 2.0 Schrödinger, LLC).

### Antibody production

To produce GldL antibodies a 462-bp fragment of *gldL* spanning the cytoplasmic domain (GldL_C_, residues 66-215) was amplified from genomic DNA using primers FL125 and FL126. This fragment was inserted between the NdeI and BamHI sites of pWALDO-sfGFPd. The resulting vector, pFL43, produces a GldL_C_-TEV-sfGFP-His_8_ fusion protein.

To produce GldM antibodies a 1305-bp fragment of *gldM* spanning the periplasmic domain (GldM_P_, residues 78-513) was amplified from genomic DNA using primers FL128 and FL129. This fragment was inserted between the NdeI and BamHI sites of pWALDO-sfGFPd. The resulting vector, pFL44, produces a GldM_P_-TEV-sfGFP-His_8_ fusion protein.

*E. coli* BL21 Star (DE3) cells containing either pFL43 or pFL44 were grown in 1.2 l LB medium at 37 °C to OD_600_ = 0.5 mid-log phase and protein expression induced by addition of 400 μM IPTG. The cells were then cultured for an additional 5 h at 20 °C. Cells were harvested by centrifugation at 6,000*g* for 25 min and stored at −20°C until further use. All purification steps were carried out at 4°C. Cell pellets were resuspended in PBS containing 30 μg ml^−1^ DNase I, 400 μg ml^−1^ lysozyme and 1 mM phenylmethylsulfonyl fluoride at a ratio of 5 ml of buffer to 1 g of cell pellet. Cells were incubated on ice for 30 min before being lysed by two passages through a TS series 1.1kW cell disruptor (Constant System Ltd) at 30,000 PSI. Cell debris was removed by centrifugation at 20,000*g* for 25 min. The supernatant was then clarified using a 0.22 μm syringe filter unit (Millipore) and circulated through a 5 ml HisTrap HP column (GE Healthcare) for 2 h. The column was washed with 10 CV of PBS containing 10 mM imidazole and bound proteins were eluted with a 10-500 mM linear imidazole in 10 CV of PBS. Peak fractions were collected, diluted with an equal volume of PBS, pH 8.0, containing 0.5 mM EDTA, and dialyzed for 1 h at 4 °C against 1 l of the same buffer. TEV-His_6_ protease was added to the pooled fractions at a 1 to 100 protein mass ratio and dialysis was continued overnight at 4°C against 1 l of PBS containing 0.5 mM EDTA and 1 mM DTT. The cleavage reaction was then circulated through a HisTrap HP column (GE Healthcare) for 2 h and the flowthrough collected. This preparation was subjected to SDS-PAGE followed by Coomassie Blue staining. The band corresponding to the recombinant protein domain was excised from the gel and used by Davids Biotechnologie GmbH (Regensburg, DE) to raise polyclonal antibodies.

### Preparation of samples for whole cell immunoblotting analysis of GldL and GldM

Strains were grown in CYE medium for 22 h, reaching OD_600_ = 5.5-6.5. The cells in 1 ml samples of the culture were harvested by centrifugation at 9,000*g* for 10 min, resuspended and washed once in 1 ml PBS, and finally resuspended in PBS to an OD_600_ = 5.0. These samples were then diluted ten-fold in PBS, subject to SDS-PAGE, and analysed by immunoblotting with GldL_C_ or GldM_P_ primary antibodies, and anti-Rabbit IgG HRP Conjugate (Promega) secondary antibodies.

### Measurement of chitinase secretion

Cells were grown in MM for 15.5 h, reaching OD_600_ = 0.75-1.25. 5 ml culture samples were subject to centrifugation at 3,720*g* for 10 min to remove cells. Chitinase activity in the resulting supernatants was determined using a fluorometric chitinase assay kit (Sigma) with the synthetic substrate 4-methylumbelliferyl N,N’-diacetyl-β-D-chitobioside. Statistical analysis of the results was carried out using GraphPad Prism version 7.03 for Windows (GraphPad Software, La Jolla California USA, www.graphpad.com). Normality of datasets was assessed by the Kolmogorov-Smirnov test. Normally distributed datasets were compared to the parental strain using one-way ANOVA followed by Dunn’s multiple comparison test. Non-normally distributed datasets were compared to WT using a Kruskal-Wallis test followed by Dunnett’s multiple comparisons test.

### Measurement of gliding motility on agar

Strains were grown overnight in PY2 medium, washed once in PY2 medium, then resuspended in Py2 medium to an OD_600_ = 0.1. A 2 μl sample was then spotted on PY2 agar plates. Plates were incubated at 25°C for 48 h before imaging with a Zeiss AXIO Zoom MRm CCD camera and Zeiss software (ZenPro 2012, version 1.1.1.0).

### Microscopic observation of gliding motility on glass

Strains were grown overnight in MM, diluted 40-fold into fresh medium and grown for 5 h at 25°C with 50 rpm shaking. An aliquot of the culture was introduced to a tunnel slide, incubated for 5 min, washed twice with 100 μl fresh medium and imaged in phase contrast on a Zeiss Axoivert 200 microscope fitted with a Photometrics Coolsnap HQ Camera using a 63x 1.4NA Plan-Apochromat lens. Tif images were captured using Metamorph software (Molecular Devices) and then modified in Image J^45^. Videos were collected at 20 frames per second with a 15 ms exposure time using 2×2 binning.

### Single molecule fluorescence imaging of SprB mobility

Strains expressing the Halo Tag-SprB fusion were grown overnight in CYE medium, diluted 40-fold into fresh medium and grown at 30°C with 180 rpm shaking to early stationary phase (OD_600_ = 0.8-1.0). The HaloTag-SprB fusion was labelled by mixing 1 ml of this culture with 1 μl of a 10 μM stock solution of PA Janelia Fluor 646 HaloTag ligand^46,47^ in dimethyl sulfoxide (DMSO) and cultured for a further 30 min. The cells were harvested at 9000*g* for 3 min, washed 3 times with 650 μl PY2 medium, and then 2 μl cells were spotted onto a 1 % agarose pad containing 50 % PY2 medium.

Fluorescence images were acquired at 25°C using a Nanoimager (Oxford Nanoimaging) equipped with 405nm and 640nm 1W DPSS lasers. Optical magnification was provided by a 100x oil-immersion objective (Olympus, numerical aperture (NA) 1.4) and images were acquired using an ORCA-Flash4.0 V3 CMOS camera (Hamamatsu). Cells were imaged using a 20 ms exposure time, with the 405 nm photoactivation laser at 10% power and the 640 nm measurement laser at 20% power. Different strains were imaged at different strobing frequencies to accommodate the large differences in adhesin velocities between the different *gldLM* mutants. Specifically, strain Ak_73 (wild type *gldLM*) was imaged without strobing, strains Ak_196 (*gldL(Y13A)*), Ak_197 (*gldL(H30A)*), and Ak_199 (*gldM(Y17F)*) were imaged with a 60 ms dark interval, whilst Ak_198 (*gldL(K27A)*) was imaged with a 480 ms dark interval.

Fluorescent foci were tracked using the Nanoimager software. Helical tracks were exported as csv files and the average frame to frame displacement calculated. The resulting data were analysed using GraphPad Prism version 7.03 for Windows (GraphPad Software, La Jolla California USA, www.graphpad.com), using a Kruskal-Wallis test followed by Dunn’s multiple comparisons test. The data were plotted using Prism.

### Pulse-chase assay of Type 9 protein export

The required *F. johnsoniae* strain was inoculated from a freshly-streaked plate into 5 ml SDY medium (SD medium^48^ containing 0.01% yeast extract) and cultured for 24 h at 30 °C with shaking. The cultures were harvested by centrifugation, resuspended in the same volume of SDAC (SD medium with 0.04 mg ml^−1^ of all L-amino acids except methionine plus 1.5% CYE medium), then diluted one in five in fresh SDAC. These cultures were grown for 2.5 h at 30 °C and 30 rpm shaking. 40 μCi ml^−1^ EasyTag L-[^35^S]-methionine (Perkin-Elmer) was then added and growth was continued for 30 min. Cultures were harvested by centrifugation and resuspended in the same volume of unlabelled SDAC containing 0.4 mg ml^−1^ unlabelled L-methionine, and where required, either 10 μM carbonyl cyanide *m*-chlorophenyl hydrazone (CCCP) or 10 mM sodium arsenate. The cultures were further incubated with shaking at 30 °C. At appropriate time points 1 ml samples were removed from each culture and the cells pelleted by centrifugation. Cells were resuspended in RIPA buffer (10 mM Tris-HCl pH 7.5, 150 mM NaCl, 0.5 mM EDTA, 0.1% SDS, 1% Triton X-100, 1% deoxycholate, 0.09% sodium azide). Both the resuspended pellets and culture supernatants stored at 4 °C until the end of the time course, at which point all samples were incubated for a further 1 hour at 4 °C. The cell pellet samples were then diluted with 1.5 volumes of RND buffer (10 mM Tris-HCl pH 7.5, 150 mM NaCl, 0.5 mM EDTA), centrifuged for 10 min at 13,000*g* and the supernatant retained. Both these cell pellet-derived samples and the culture supernatant samples were incubated with 10 μl RFP-trap MA (Chromotek) for one hour at 24 °C with constant mixing. The RFP-trap resin was isolated on a magnetic rack and washed with 2 × 800 μl RND buffer, then resuspended in SDS-PAGE sample buffer. Samples were separated by SDS-PAGE, and the resulting gels fixed, dried and exposed to radiographic film (GE Healthcare).

### Measurement of cellular ATP levels

Strains were cultured as for the pulse-chase experiments to the point at which the cells were transferred to SDAC medium. The cells were then incubated for 3 h at 30 °C and 110 rpm. 0.5 ml samples were treated with 10 μM CCCP, or 10 mM sodium arsenate, or left untreated for 20 min at 24 °C. Cellular ATP levels were then determined using an ATP Bioluminescence Assay Kit HS II (Roche). Cells were diluted to OD_600_ = 0.1 with dilution buffer from the kit. 200 μl aliquots of diluted cells were combined with 200 μl lysis buffer from the kit and incubated at 70 °C for 5 min. Lysates were immediately transferred to ice and then clarified by centrifugation. The ATP content in the lysates was determined according to the kit manufacturer’s instructions and using a CLARIOstar Plus plate reader (BMG Labtech). Statistical analysis of cellular ATP levels was carried out using R ^49^.

## Data availability

The cryo-EM volumes have been deposited in the Electron Microscopy Data Bank (EMDB) with accession codes EMD-10893 and EMD-10894, and the coordinates have been deposited in the Protein Data Bank (PDB) with accession code 6YS8. Source Data are available with the online version of the paper.

## Extended data figures and tables

**Extended Data Fig. 1.**
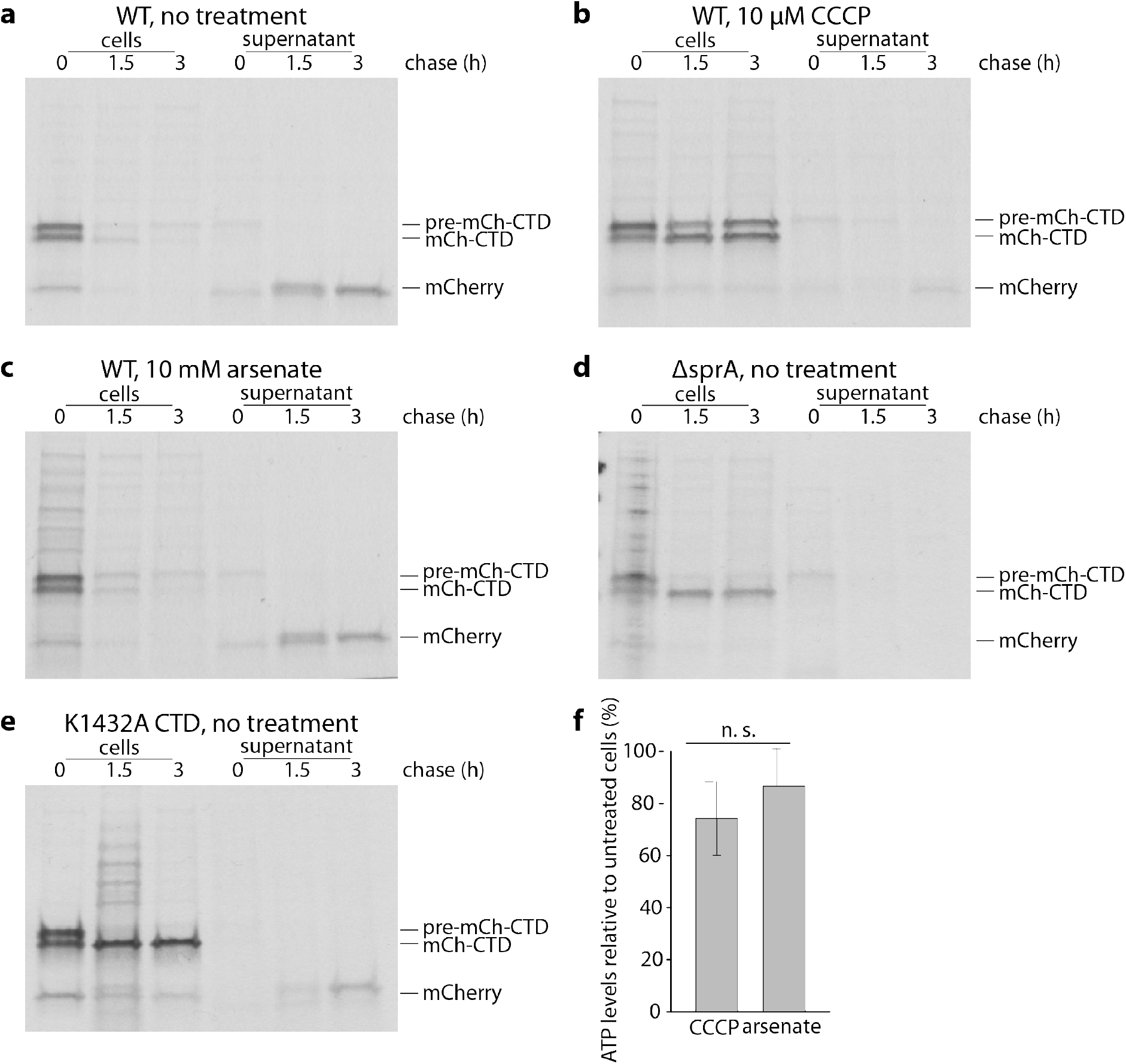
Protein export by the T9SS requires the protonmotive force. **a-e,** Pulse-chase analysis of the export of a signal sequence-mCherry-CTD (T9SS-targeting C-terminal domain) fusion protein by the *F. johnsoniae* T9SS. Cells were labelled with [^35^S]methionine for 30 min, chased with cold methionine for 0 to 3 h (as indicated), and then separated into cell and supernatant (culture medium) fractions. Fusion protein was enriched by anti-mCherry immunoprecipitation and then analysed by SDS-PAGE and autoradiography. Similar data were obtained for two biological repeats. Pre-mCh-CTD, full length fusion protein; mCh-CTD, fusion protein from which the signal sequence has been removed; mCh, mCherry from which both the signal sequence and CTD have been removed. Export of the fusion protein was blocked by treatment with the protonophore carbonyl cyanide m-chlorophenyl hydrazone (CCCP, panel **b**) but not by the ATP synthase inhibitor arsenate (panel, **c**). Control experiments confirm that the observed export of the mCherry fusion requires the T9SS (Δ*sprA* strain Fl_004 ^20^, panel **d**) and a functional CTD (non-functional K1432A CTD^50^, panel **e**). Note, that transport of the fusion across the cytoplasmic membrane by the Sec apparatus is also inhibited by CCCP (panel **c**) in agreement with previous observations in *Escherichia coli*^51^. **f,** Measurements of whole cell ATP levels confirm that the effects of CCCP on protein transport are not an indirect effect of decreased cellular ATP. Bioluminescence readings from treated cells were normalised to those of untreated cells from the same starting culture. CCCP and arsenate concentrations were as in panels **b**) and **c**). Error bars represent 1 SD (10 biological repeats). The datasets were tested for normality using the Shapiro-Wilk test and also inspected visually using Q-Q plots. The Bartlett test was used to establish that the variances of the two datasets were likely homogeneous. An independent two-tailed t-test indicated that the ATP levels in CCCP- and arsenate-treated cells were not significantly different (n.s.) (t-value = −1.9619, df = 17.994, p-value = 0.06543>0.05) even though only CCCP treatment blocks T9SS protein export.

**Extended Data Fig. 2.**
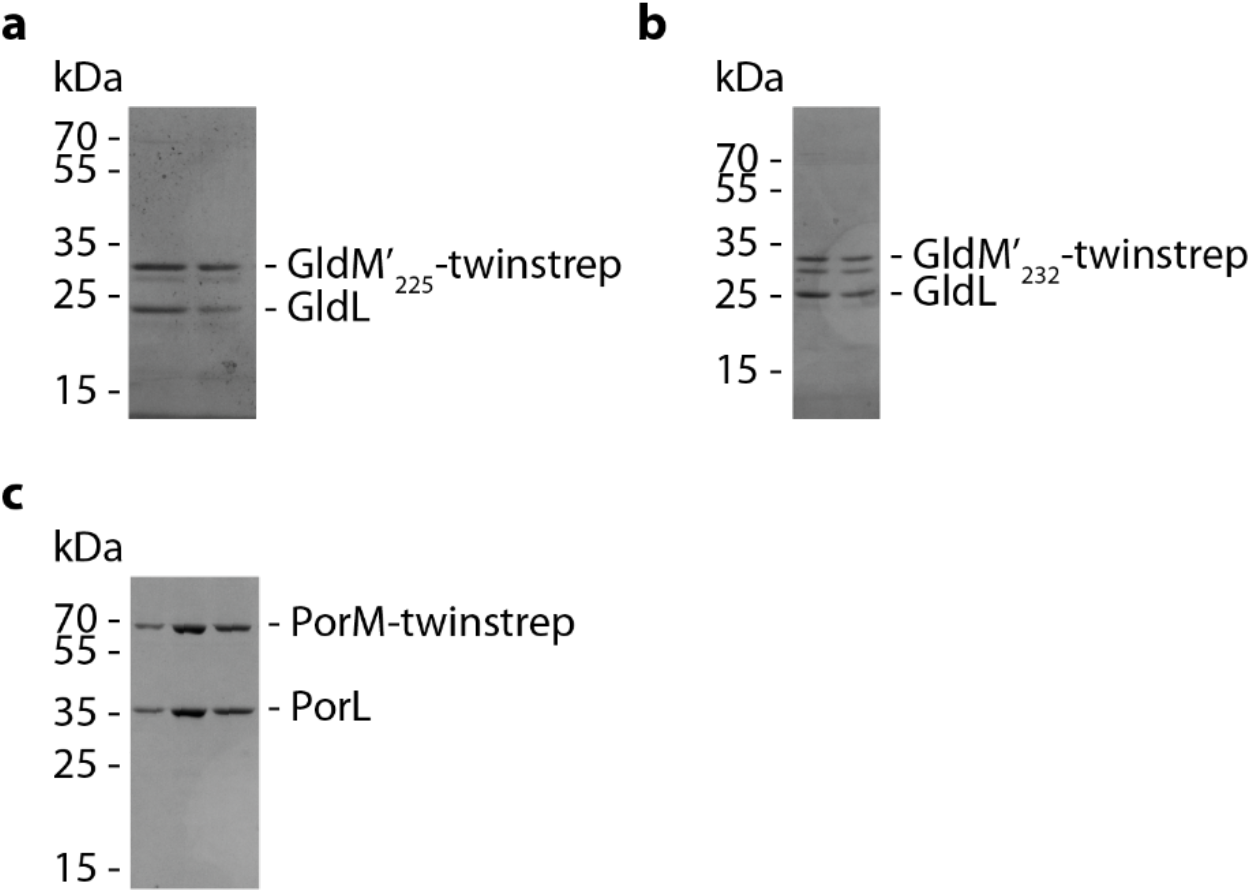
Purification of the GldLM and PorLM complexes. Coomassie Blue-stained SDS-PAGE gels of the (**a**) GldLM_225_(produced from pRHJ007), (**b**) GldLM_232_ (produced from pRHJ008), and (**c**) PorLM (produced from pRHJ006) complexes used for cryo-EM structure determination. In each case the size exclusion column fractions that were pooled to make the cryo-EM grids are shown.

**Extended Data Fig. 3.**
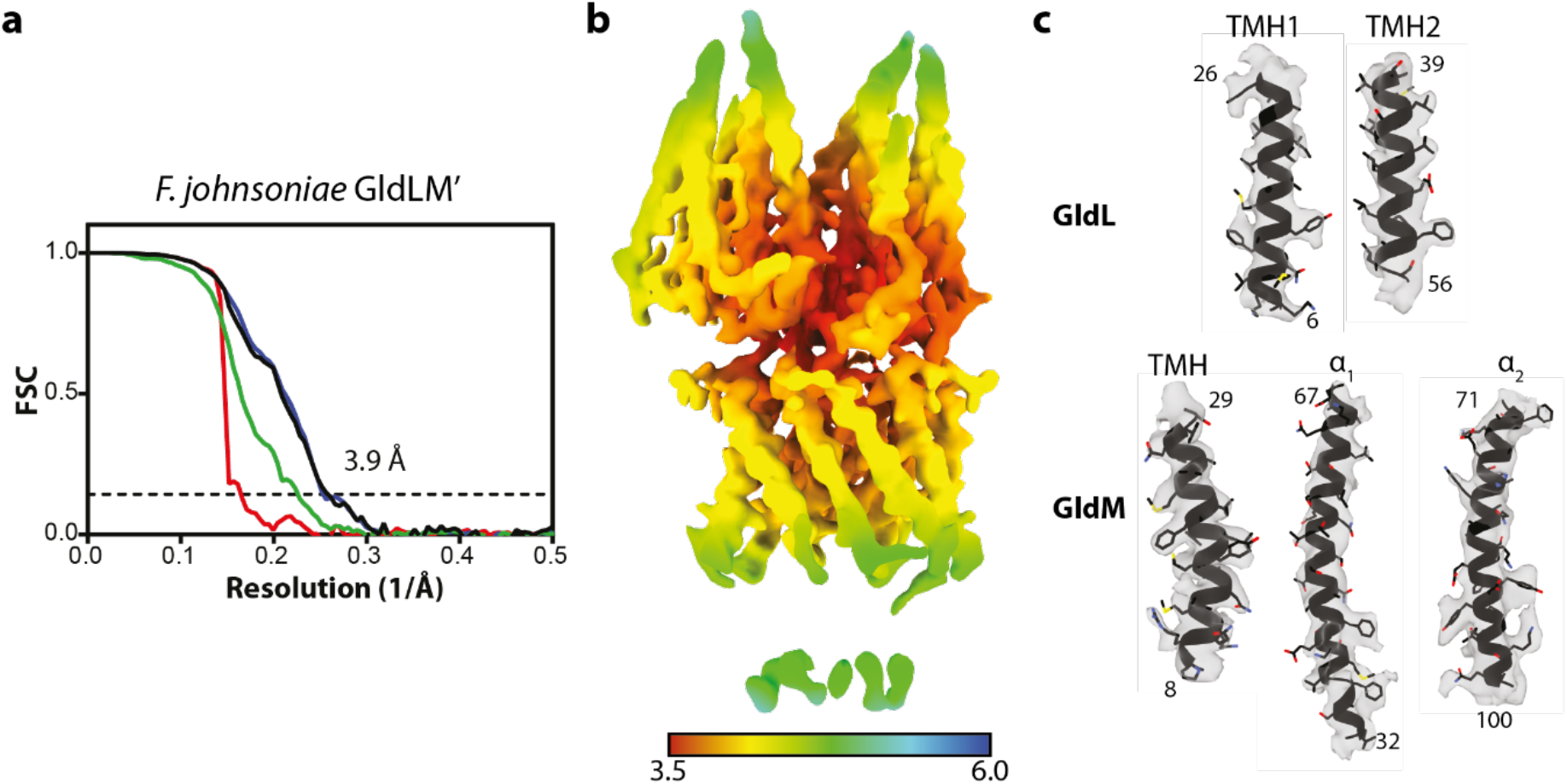
Cryo-EM map quality and resolution estimates. **a** Fourier Shell Correlation (FSC) plot for the GldLM’ structure. The resolution at the gold-standard cutoff (FSC = 0.143) is indicated by the dashed line. Curves: Red, phase-randomized; Green, unmasked; blue, masked; black, MTF-corrected **b** Local resolution estimates (in Å) for the sharpened GldLM’ map. **c** Representative modelled densities.

**Extended Data Fig. 4.**
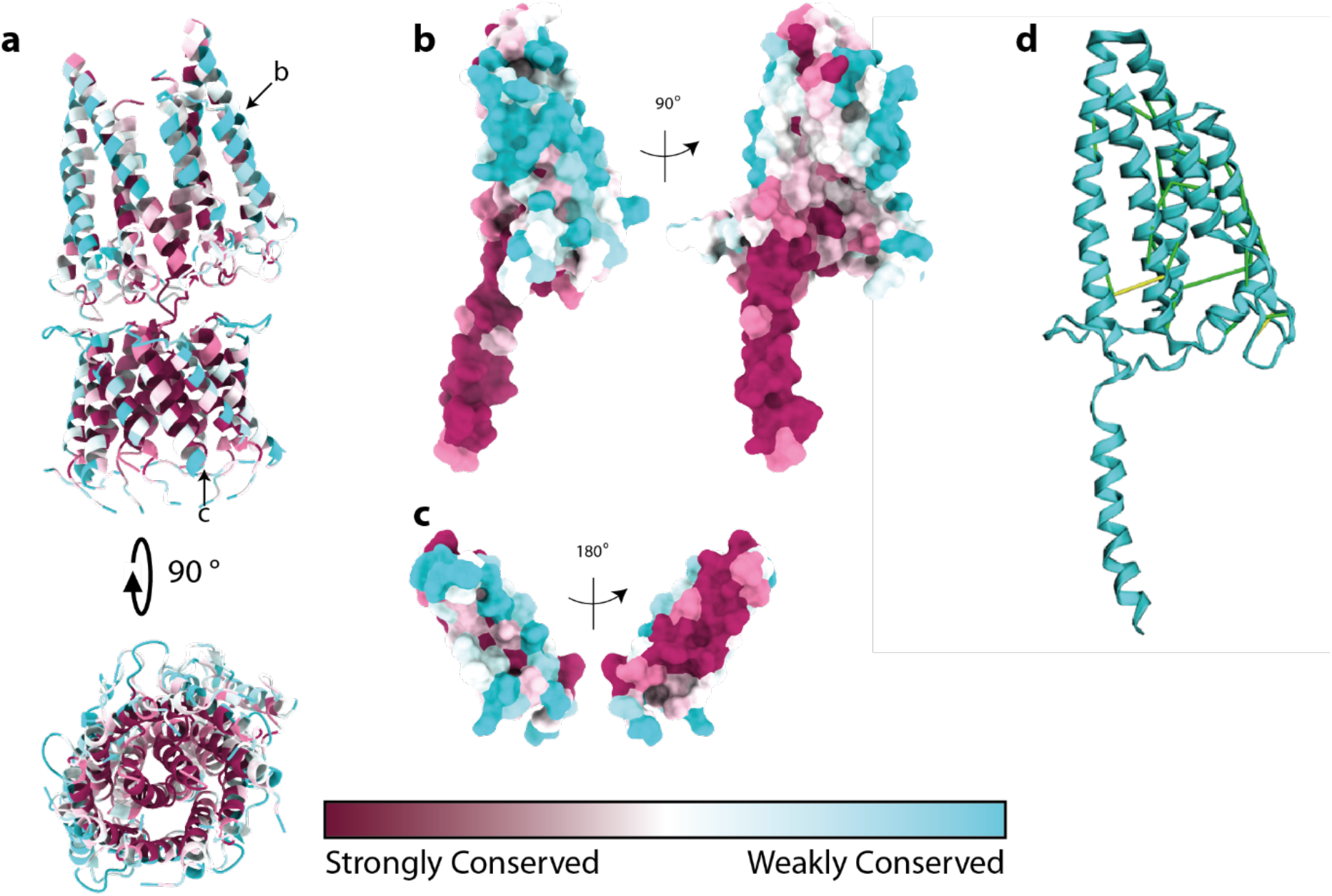
Conservation analysis of the GldLM complex. **a-c,** Sequence conservation assessed using Consurf^43^. **a,** The whole complex in cartoon representation. The chains shown individually in **b** and **c** are indicated with black arrows. **b,** GldM (chain B) and **c,** GldL (chain F) in surface representation. The left hand panels show the chains in the same orientation as the upper panel of **a**. **d,** Covarying residue pairs in the first periplasmic domain of GldM identified using the Gremlin server ^52^. The highest-scoring pairs (score cut-off 1.63) are shown on the structure of the GldLM’ complex presented here. Contacts with a minimum atom-atom distance of <5 Å are shown in green, and 5-10 Å in yellow. Note, that no high-scoring intermolecular pairs were observed.

**Extended Data Fig. 5.**
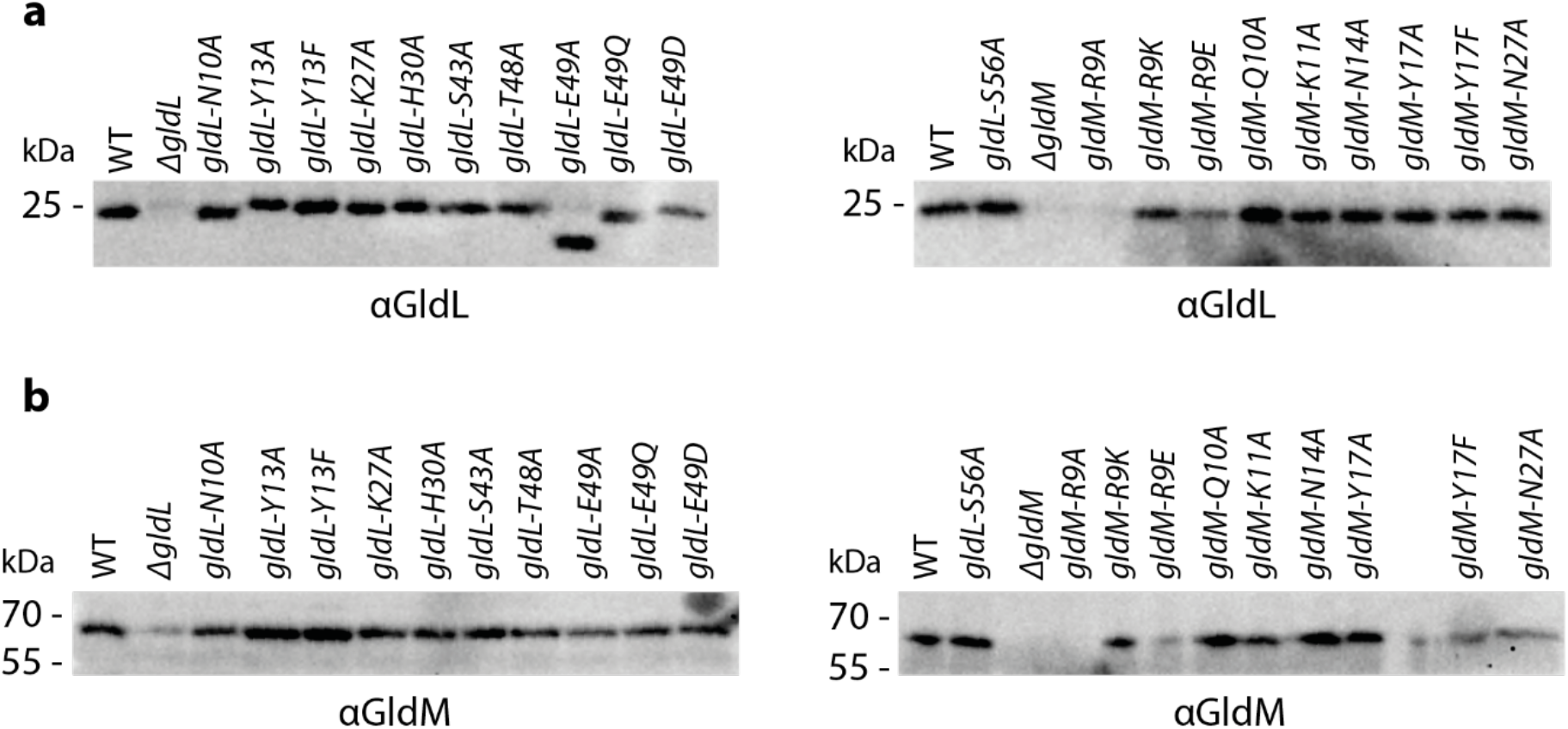
GldL and GldM expression levels in *gldL* and *gldM* mutant strains. Whole cell immunoblotting of the indicated strains for the presence of (**a**) GldL and (**b**) GldM. Similar data were obtained for three independent preparations.

**Extended Data Table 1.** Cryo-EM data collection, refinement and validation statistics.

|  | F. johnsoniae GldLM<br>(EMD-10893)<br>(PDB 6YS8) | P. gingivalis PorLM<br>(EMD-10894) |
| --- | --- | --- |
| **Data collection and processing** |  |  |
| Magnification | 165,000 | 165,000 |
| Voltage (kV) | 300 | 300 |
| Electron exposure (e-/Å<sup>2</sup>) | 48 | 48 |
| Defocus range (µm) | 1.0-3.0 | 1.0-3.0 |
| Pixel size (Å) | 0.822 | 0.822 |
| Symmetry imposed | C1 | C1 |
| Initial particle images (no.) | 3,284,887 | 1,133,336 |
| Final particle images (no.) | 119,230 | 199,929 |
| Map resolution (Å) | 3.9 | 8.6 |
| FSC threshold | 0.143 | 0.143 |
| Map resolution range (Å) | 3.5-5.8 |  |
| **Refinement** |  |  |
| Initial model used (PDB code) | None |  |
| Model resolution (Å) | 3.9 |  |
| FSC threshold | 0.143 |  |
| Model resolution range (Å) | 3.5-5.8 |  |
| Map sharpening B factor (Å<sup>2</sup>) | -131 |  |
| Model composition |  |  |
| Non-hydrogen atoms | 5624 |  |
| Protein residues | 737 |  |
| Ligands | 0 |  |
| B factors (Å<sup>2</sup>) |  |  |
| Protein | 117 |  |
| Ligand | NA |  |
| R.m.s. deviations |  |  |
| Bond lengths (Å) | 0.007 |  |
| Bond angles (°) | 1.186 |  |
| Validation |  |  |
| MolProbity score | 2.22 |  |
| Clashscore | 10.52 |  |
| Poor rotamers (%) | 1.70 |  |
| Ramachandran plot |  |  |
| Favored (%) | 91.42 |  |
| Allowed (%) | 6.36 |  |
| Disallowed (%) | 2.21 |  |

**Extended Data Table 2.** Contacts between the periplasmic loops of GldL and GldM. The PDBePISA server (http://www.ebi.ac.uk/pdbe/pisart.html)^53^was used to calculate the sizes of the interfaces between the periplasmic loops of each copy of GldL and the periplasmic domains of GldM. The periplasmic loops of GldL were taken to include residues 29-41.

| GldM Copy | GldL Copy | GldL Buried Surface area (Å<sup>2</sup>) | Number of GldL Residues | GldL Residues | GldM Residues |
| --- | --- | --- | --- | --- | --- |
| A | C | 265.4 | 8 | Thr29, His30, Phe31, Glu32, Thr37, Gly38, Thr39, Val40 | Arg120, Asp122, Asp125, Asp126, Phe128, Thr129, Gly130, Lys193 |
| A | D | 107.4 | 3 | Thr29, His30, Thr39 | Met26, Lys30, Glu116, Ala117, Asp119, Arg120 |
| A | E | 139.5 | 4 | Thr29, His30, Phe31, Glu32 | Met26, Asn27, Ser29, Lys30, Glu31 |
| A | F | 125.8 | 3 | His30, Thr37, Thr39 | Lys180, Glu181, Ile183 |
| A | G | 183.8 | 4 | Thr29, His30, Thr37, Thr39 | Thr129, Gly130, Asp131, Asn179, Ile183, Lys184, Glu185 |
| B | C | - | - | - | - |
| B | D | - | - | - | - |
| B | E | 64.4 | 5 | Phe31, Glu32, Thr37, Thr39, Val40 | Arg120, Asp122, Lys180, Gly194 |
| B | F | 97.2 | 2 | Thr29, His30 | Val28, Ser29, Ile33, Glu116, Ala117, Asp119 |
| B | G | 18.1 | 1 | His30 | Met26 |

**Extended Data Table 3.** Gliding motility and T9SS phenotypes of mutant strains.

| Strain | Gliding on PY2 agar | Gliding on glass | Chitinase assay (arbitrary units ± s.d.) | SprB propulsion (nm s<sup>-1</sup> ± s.d.) | GldL stability | GldM stability |
| --- | --- | --- | --- | --- | --- | --- |
| WT | +++ | Glides normally | 1.00 ± 0.2 | 1358 ± 900 | Present | Present |
| Δ*gldL* | - | Does not adhere | 0.00 ± 0.0 | n.d. | None | Reduced |
| *gldL*<sub>N10A</sub> | +++ | Glides normally | 1.09 ± 0.3 | n.d. | Present | Present |
| *gldL*<sub>Y13A</sub> | - | Adheres, does not glide | 0.41 ± 0.1 | 275 ± 100 | Present | Present |
| *gldL*<sub>Y13F</sub> | - | Adheres, does not glide | 1.37 ± 0.5 | n.d. | Present | Present |
| *gldL*<sub>K27A</sub> | - | Adheres, does not glide | 0.40 ± 0.1 | 31 ± 20 | Present | Present |
| *gldL*<sub>H30A</sub> | ++ | Glides slowly | 1.16 ± 0.2 | 524 ± 200 | Present | Present |
| *gldL*<sub>S43A</sub> | +++ | Glides normally | 0.96 ± 0.3 | n.d. | Present | Present |
| *gldL*<sub>T48A</sub> | +++ | Glides normally | 1.15 ± 0.6 | n.d. | Present | Present |
| *gldL*<sub>E49A</sub> | - | Does not adhere | 0.00 ± 0.0 | n.d. | Clipped | Present |
| *gldL*<sub>E49Q</sub> | - | Does not adhere | 0.00 ± 0.0 | n.d. | Present | Present |
| *gldL*<sub>E49D</sub> | - | Very few cells adhere | 0.02 ± 0.0 | n.d. | Present | Present |
| *gldL*<sub>S56A</sub> | +++ | Glides normally | 0.88 ± 0.2 | n.d. | Present | Present |
| Δ*gldM* | - | Does not adhere | 0.00 ± 0.0 | n.d. | None | None |
| *gldM*<sub>R9A</sub> | - | Does not adhere | 0.00 ± 0.0 | n.d. | None | None |
| *gldM*<sub>R9K</sub> | ++ | Glides slowly | 0.95 ± 0.2 | 486 ± 200 | Present | Present |
| *gldM*<sub>R9E</sub> | - | Does not adhere | 0.01 ± 0.0 | n.d. | Reduced | Reduced |
| *gldM*<sub>Q10A</sub> | +++ | Glides normally | 0.98 ± 0.3 | n.d. | Present | Present |
| *gldM*<sub>K11A</sub> | +++ | Glides normally | 0.91 ± 0.1 | n.d. | Present | Present |
| *gldM*<sub>N14A</sub> | +++ | Glides normally | 1.10 ± 0.3 | n.d. | Present | Present |
| *gldM*<sub>Y17A</sub> | - | Adheres, does not glide | 0.11 ± 0.0 | n.d. | Present | Present |
| *gldM*<sub>Y17F</sub> | + | Glides slowly | 1.13 ± 0.4 | 774 ± 400 | Present | Present |
| *gldM*<sub>N27A</sub> | +++ | Glides normally | 1.30 ± 0.5 | n.d. | Present | Present |

**Extended Data Table 4.** Bacterial strains used in this study.

| Strain | Genotype | Reference |
| --- | --- | --- |
| ***E. coli*** |  |  |
| S17-1 | *pro*, *res<sup>-</sup> hsdR17 (rK<sup>-</sup> mK<sup>+</sup>) recA<sup>-</sup>*, *RP4-2-Tc::Mu-Km::Tn7*, *Tp<sup>r</sup>* | 32 |
| BL21 Star™ (DE3) | *F- ompT hsdS<sub>B</sub> (r<sub>B</sub><sup>-</sup>, m<sub>B</sub><sup>-</sup>) gal dcm rne131 (DE3)* | Invitrogen |
| ***F. johnsoniae*** |  |  |
| UW101 |  | 54 |
| Fl_004 | UW101 Δ*sprA* | 20 |
| Fl_030 | UW101 Δ*porV* | 20 |
| Fl_082 | UW101 Δ*gldL* | This study |
| Rhj_006 | UW101 Δ*gldM* | This study |
| Rhj_004 | UW101 *gldL*-*twinstrep* | This study |
| Rhj_017 | UW101 *gldL*<sub>N10A</sub> | This study |
| Rhj_018 | UW101 *gldL*<sub>Y13A</sub> | This study |
| Rhj_024 | UW101 *gldL*<sub>Y13F</sub> | This study |
| Rhj_019 | UW101 *gldL*<sub>K27A</sub> | This study |
| Rhj_011 | UW101 *gldL*<sub>H30A</sub> | This study |
| Rhj_013 | UW101 *gldL*<sub>S43A</sub> | This study |
| Rhj_020 | UW101 *gldL*<sub>T48A</sub> | This study |
| Rhj_021 | UW101 *gldL*<sub>E49A</sub> | This study |
| Rhj_022 | UW101 *gldL*<sub>E49Q</sub> | This study |
| Rhj_025 | UW101 *gldL*<sub>E49D</sub> | This study |
| Rhj_023 | UW101 *gldL*<sub>S56A</sub> | This study |
| Rhj_035 | UW101 *gldM*<sub>R9A</sub> | This study |
| Rhj_032 | UW101 *gldM*<sub>R9K</sub> | This study |
| Rhj_029 | UW101 *gldM*<sub>R9E</sub> | This study |
| Rhj_012 | UW101 *gldM*<sub>Q10A</sub> | This study |
| Rhj_030 | UW101 *gldM*<sub>K11A</sub> | This study |
| Rhj_016 | UW101 *gldM*<sub>N14A</sub> | This study |
| Rhj_014 | UW101 *gldM*<sub>Y17A</sub> | This study |
| Rhj_015 | UW101 *gldM*<sub>Y17F</sub> | This study |
| Rhj_028 | UW101 *gldM*<sub>N27A</sub> | This study |
| Rhj_031 | UW101 *gldL*<sub>N10A</sub>-*twinstrep* | This study |
| Rhj_033 | UW101 *gldL*<sub>Y13A</sub>-*twinstrep* | This study |
| Rhj_041 | UW101 *gldL*<sub>Y13F</sub>-*twinstrep* | This study |
| Rhj_053 | UW101 *gldL*<sub>K27A</sub>-*twinstrep* | This study |
| Rhj_027 | UW101 *gldL*<sub>H30A</sub>-*twinstrep* | This study |
| Rhj_042 | UW101 *gldL*<sub>S43A</sub>-*twinstrep* | This study |
| Rhj_034 | UW101 *gldL*<sub>T48A</sub>-*twinstrep* | This study |
| Rhj_039 | UW101 *gldL*<sub>E49A</sub>-*twinstrep* | This study |
| Rhj_026 | UW101 *gldL*<sub>E49Q</sub>-*twinstrep* | This study |
| Rhj_050 | UW101 *gldL*<sub>E49D</sub>-*twinstrep* | This study |
| Rhj_038 | UW101 *gldL*<sub>S56A</sub>-*twinstrep* | This study |
| Rhj_049 | UW101 *gldL*-*twinstrep gldM*<sub>R9A</sub> | This study |
| Rhj_045 | UW101 *gldL*-*twinstrep gldM*<sub>R9K</sub> | This study |
| Rhj_040 | UW101 *gldL*-*twinstrep gldM*<sub>R9E</sub> | This study |
| Rhj_046 | UW101 *gldL*-*twinstrep gldM*<sub>Q10A</sub> | This study |
| Rhj_044 | UW101 *gldL*-*twinstrep gldM*<sub>K11A</sub> | This study |
| Rhj_051 | UW101 *gldL*-*twinstrep gldM*<sub>N14A</sub> | This study |
| Rhj_43 | UW101 *gldL*-*twinstrep gldM*<sub>Y17A</sub> | This study |
| Rhj_37 | UW101 *gldL*-*twinstrep gldM*<sub>Y17F</sub> | This study |
| Rhj_052 | UW101 *gldL*-*twinstrep gldM*<sub>N27A</sub> | This study |
| Ak_73 | UW101 Δ*porV* *halotag::sprB* | This study |
| Ak_203 | AK_73 Δ*gldM* | This study |
| Ak_205 | AK_73 Δ*gldL* | This study |
| Ak_196 | AK_73 *gldL*<sub>Y13A</sub> | This study |
| Ak_197 | AK_73 *gldL*<sub>H30A</sub> | This study |
| Ak_198 | AK_73 *gldL*<sub>K27A</sub> | This study |
| Ak_199 | AK_73 *gldM*<sub>Y17F</sub> | This study |
| Ak_289 | AK_73 *gldM*<sub>R9K</sub> | This study |

**Extended Data Table 5.**
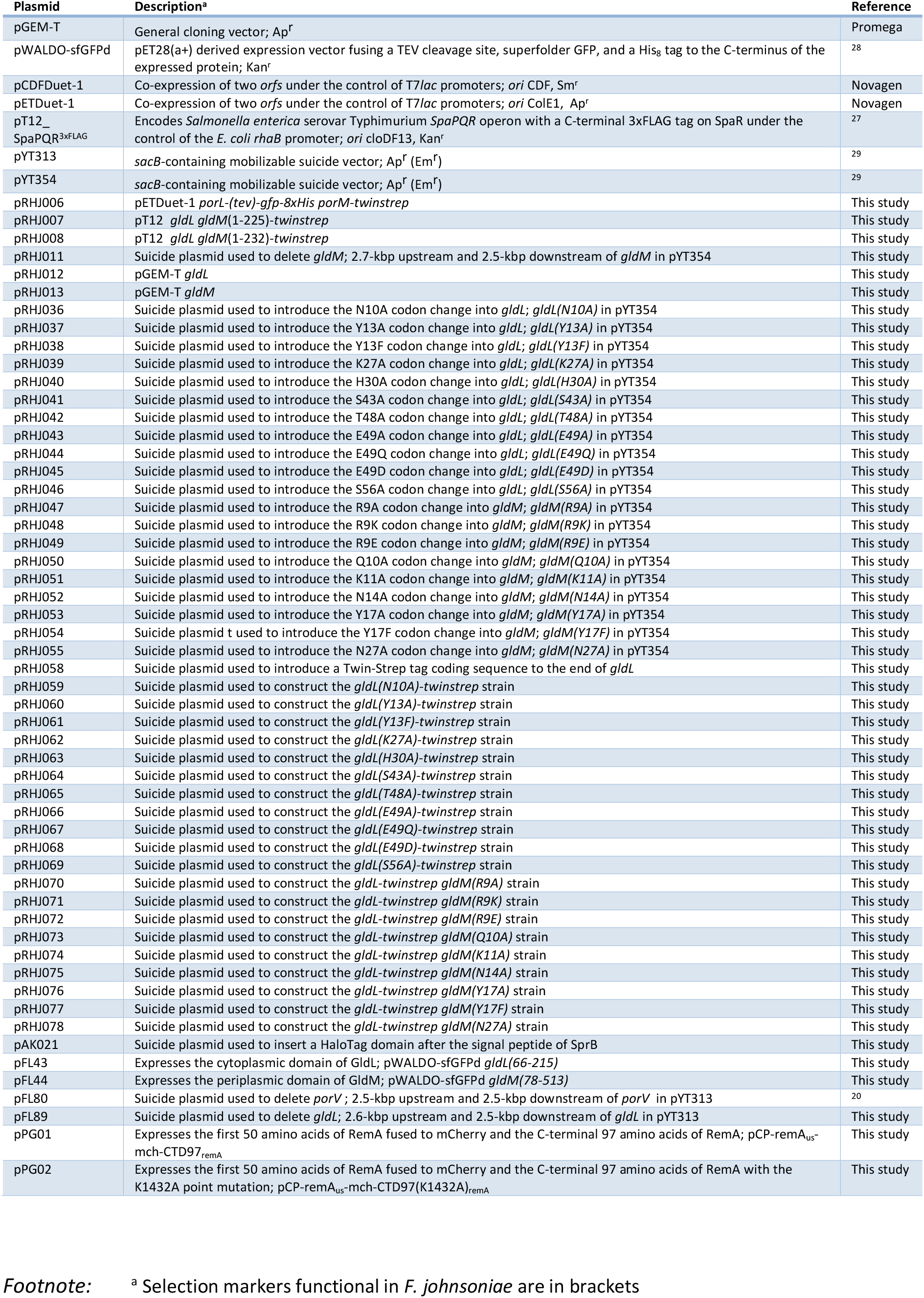
Plasmids used in this study.

**Supplementary Data Table 1:** Oligonucleotides used in this study.

**Supplementary Data Video1:** Mutant strains gliding on glass. WT, *gldL*_*Y13A*_, *gldL*_*Y13F*_, *gldL*_*K27A*_, *gldL*_*H30A*_, *gldL*_*E49D*_, *gldM*_*R9K*_, *gldM*_*Y17A*_ and *gldM*_*Y17F*_ cells are shown.

**Supplementary Data Video 2:** Fluorophore-labelled SprB adhesin moving on the surface of a WT cell.

**Supplementary Data Video 3:** Fluorophore-labelled SprB adhesin moving on the surface of a *gldL*_*Y13A*_ cell.

**Supplementary Data Video 4:** Fluorophore-labelled SprB adhesin moving on the surface of a *gldL*_*K27A*_ cell.

**Supplementary Data Video 5:** Fluorophore-labelled SprB adhesin moving on the surface of a *gldL*_*H30A*_ cell.

**Supplementary Data Video 6:** Fluorophore-labelled SprB adhesin moving on the surface of a *gldM*_*R9K*_ cell.

**Supplementary Data Video 7:** Fluorophore-labelled SprB adhesin moving on the surface of a - *gldM*_*Y17F*_ cell.

**Supplementary Data Video 8:** Animation of a rotary model of GldLM function.

## Notes

### Competing Interest Statement

The authors have declared no competing interest.

